# The binding of palonosetron and other antiemetic drugs to the serotonin 5-HT3 receptor

**DOI:** 10.1101/2020.02.14.947937

**Authors:** Eleftherios Zarkadas, Hong Zhang, Wensheng Cai, Gregory Effantin, Jonathan Perot, Jacques Neyton, Christophe Chipot, Guy Schoehn, Francois Dehez, Hugues Nury

## Abstract

Inaccurately perceived as niche drugs, antiemetics are key elements of cancer treatment alleviating the most dreaded side effect of chemotherapy. Serotonin 5-HT3 receptor antagonists are the most commonly prescribed class of drugs to control chemotherapy-induced nausea and vomiting (CINV). These antagonists have been clinically successful drugs since the 1980s, yet our understanding of how they operate at the molecular level has been hampered by the difficulty of obtaining structures of drug-receptor complexes. Here, we report the cryo-EM structure of the palonosetron-bound 5-HT3 receptor. We investigate the binding of palonosetron, granisetron, dolasetron, ondansetron, and cilansetron using molecular dynamics, covering the whole set of antagonists used in the clinical practice. The structural and computational results yield detailed atomic insight into the binding modes of the drugs. In light of our data, we establish a comprehensive framework underlying the inhibition mechanism by the -setron drug family.

## Introduction

In the realm of serotonin receptors, the 5-HT3 receptor stands apart as the only ion channel^1^. It belongs to the family of pentameric ligand-gated ion channels (pLGICs)^2^. The pLGIC family comprises, in mammals, members activated by serotonin, acetylcholine, glycine and γ-aminobutyric acid. They share a common architecture: neurotransmitters bind to extracellular interfaces between subunits; the transmembrane domain (TMD) possesses 5×4 helices and features a central gated pore; the dispensable intracellular domain (ICD) is for a good part intrinsically disordered.

The interest in 5-HT3 receptor pharmacology surged when chemotherapies became first available (e.g. cisplatin approved in 1978 for ovarian and testicular cancers). At the time of their discovery in the 1980s, the first competitive antagonists were characterized as the *“most potent drugs of any pharmacological class so far described”*^3^, and less than a decade later the antiemetic research on 5-HT3 receptors was considered done by the pharma industry with numerous clinically and commercially successful drugs (coined-setrons) available^4^. Beyond emesis, fundamental and pre-clinical research indicate that the 5-HT3 receptor plays a role in depression^5^ and in body weight control^6,7^; alosetron is used clinically to treat irritable bowel syndrome^8^, vortioxetine is an antidepressant non-selectively targeting the serotonin transporter and receptors^9^.

The molecular mode of action of antiemetics has been explored extensively through binding, mutagenesis and electrophysiology studies, while structural studies on soluble model proteins gave approximate poses for the bound drugs^10,11^. Recently cryo-EM structures of the 5-HT3 receptor have revealed the conformations of the apo state, of the tropisetron-bound inhibited and of two serotonin bound states^12–14^. However, the limited resolution of the tropisetron-bound structure precluded a thorough analysis of its binding.

Here, we report a new cryo-EM structure of the 5-HT3 receptor in its inhibited state, binding palonosetron, and solved at a 2.8-Å resolution, germane for structural pharmacology. Palonosetron possesses the highest affinity in the -setron family, a slow dissociation rate and nanomolar IC50s (∼0.2 nM when pre-incubated, ∼80 nM when co-applied)^15^. Leaning on this structure, and that recently reported for the granisetron-bound 5-HT3 receptor^16^, we set out to provide a comprehensive structural framework that established the mode of action of antiemetics. Our palonosetron-bound structure served as a template for molecular dynamics (MD) simulations not only with palonosetron, but also with cilansetron, dolasetron, granisetron and ondansetron to elucidate the general rules that underlie binding of the setron family to the 5-HT3 receptor. We further establish a relationship between the body of biochemical and pharmacological data available in light of our findings.

## Results and discussion

### Global conformation

Cryo-EM imaging of the mouse homomeric 5-HT3A receptor, in the presence of palonosetron, yielded a 2.8 Å reconstruction with well-defined densities throughout the three domains of the protein (Figure 1 and SI1) - except at the level of the intrinsically disordered region of the intracellular domain (residues 329 to 399, see Figure SI2 for topology and numbering), for which the density is averaged out. The global conformation corresponds to that of previously determined inhibited states (4PIR by a nanobody form^17^, 6HIS by tropisetron^13^, 6NP0 by granisetron^16^). Compared with the previous structures, this one and the granisetron-bound one stand out in quality and may be used as templates in future modeling work, where the inhibited state is relevant. Non-proteinic densities in the orthosteric sites could be attributed non-ambiguously to palonosetron molecules (Figure 2b).

**Figure 1.**
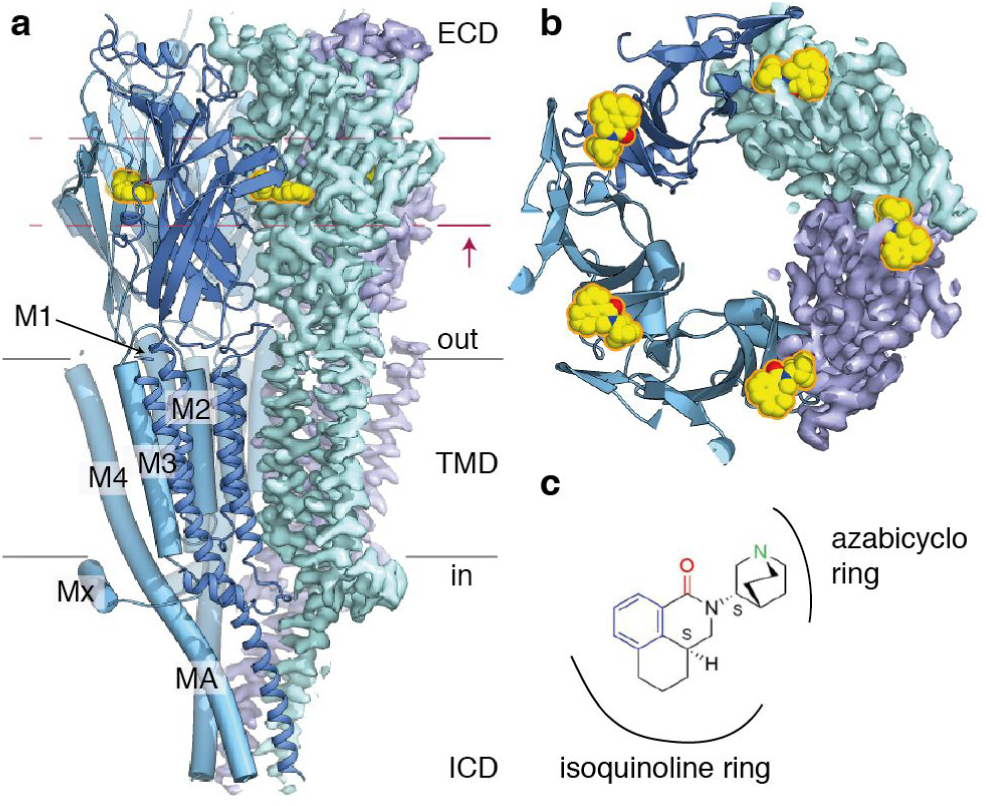
Cryo-EM structure of the mouse 5-HT3 receptor in complex with palonosetron. **a**. View parallel to the membrane plane. Three subunits are represented as cartoons while the cryo-EM map is depicted for the two last subunits. Palonosetron molecules are depicted as yellow spheres. **b**. View of a slab (fuschia lines) perpendicular to the membrane (as indicated by the arrow). **c**. Chemical formula of palonosetron, with its aromatic moiety in blue, its ring-embedded protonated nitrogen in green and its H-bond acceptor oxygen group in red. The ‘s’ letters next to chiral carbon atoms indicate stereochemistry.

**Figure 2.**
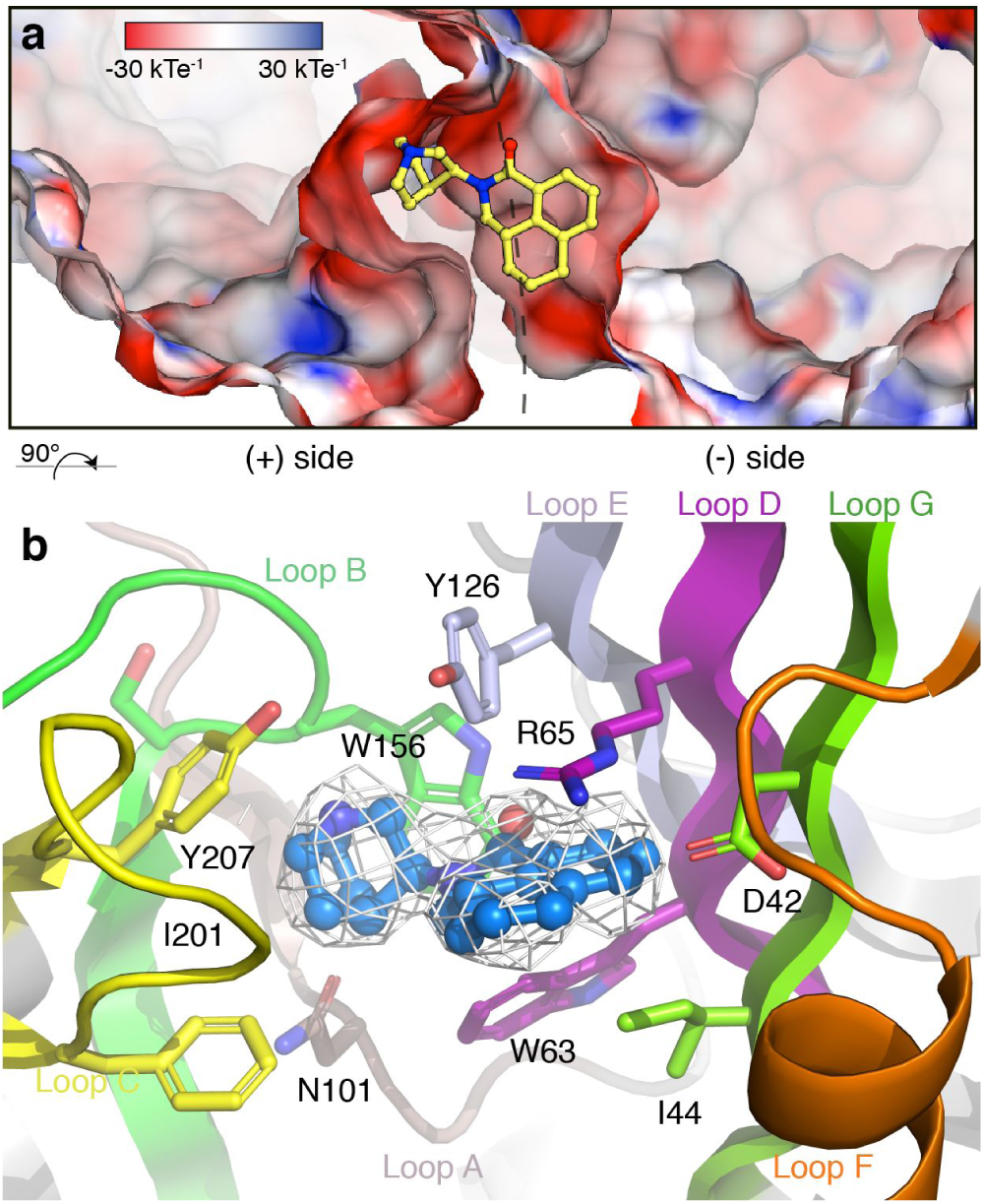
Close-up of the palonosetron binding site. **a**. Electrostatic potential mapped on the molecular surface of the 5-HT3 receptor, with stick representation of palonosetron in the neurotransmitter binding site. The view is from outside, approximately perpendicular to the membrane plane. **b**. Cartoon and stick representation, approximately parallel to the membrane. Side chains within 4 Å of the drug are depicted as sticks. The protein is colored using the same code throughout the report (on the principal side, loop A: maroon, loop B: green, loop C: yellow; on the complementary side, loop D: purple, loop E: orchid, loop F: orange, loop G: forest green). The cryo-EM density for palonosetron is depicted as a grey mesh.

### General features of the -setron binding site

Palonosetron binds tightly in the classical neurotransmitter site at the subunit/subunit interface (Figure 1). The occluded surface area between two subunits, for the extracellular domain (ECD) only, amounts to ∼1400 Å^2^. Palonosetron acts as a molecular glue, each molecule locking the conformation of the interface and shielding an extra ∼170 Å^2^ of the principal subunit and ∼210 Å^2^ of the complementary subunit from the solvent. MD simulations essentially provide the same picture (Figure SI3). The conformational dynamics of the drugs within their binding sites induce variability in the shielding of the principal subunit, concomitant with the dynamics of the H-bond network (see below).

Like every -setron drug, palonosetron possesses two moieties (Table SI1). The azabicyclo ring penetrates deep into the binding pocket, while the isoquinoline ring sits in a cavity at the surface of the complementary subunit, capped by loop C of the principal subunit (Figure 2b). The drug remains partly solvent-accessible between loop C and loop F (and one could imagine derivatives with probes grafted there).

Four aromatic residues (W156, Y126, Y207, W63) forming the so-called ‘aromatic cage’ tightly wrap around the azabicyclo ring (Figure 2b, 3a). Mutants harboring non-aromatic side chains at any of these four positions show either much decreased inhibition by palonosetron, or simply no response to serotonin^18^. Even conservative mutations such as W156Y^1^ or Y126F result in a 180- and 20-fold loss of inhibition, respectively^18^.

The palonosetron isoquinoline moiety is constrained by hydrophobic interactions with Y126, W63, I44 and I201 (Figure 2b, 3a). It also interacts with R65, the conformation of which is particularly interesting: the arginine extends over palonosetron, possibly engaging in a cation-π interaction, as reported for granisetron^10^ (the conformation of loop D residues is further discussed in Figure SI4).

A dense network of interactions stabilizes the side chains of Y126, R65, Y67, D42, R169, D177 on the complementary side (Figure 3 and Figure SI5). During simulations of all drugs, the D42-R65 and D42-R169 salt bridges are always established, whereas the D177-R169 one is occasionally disrupted. While the side-chain conformations in the binding loops of the complementary subunit strikingly resemble those of the serotonin-bound receptor, it is worth noting that both D42-R65 and D177-R169 (to a lesser extent) can break in simulations of the apo or serotonin-bound receptor (Figure SI5e,f).

**Figure 3.**
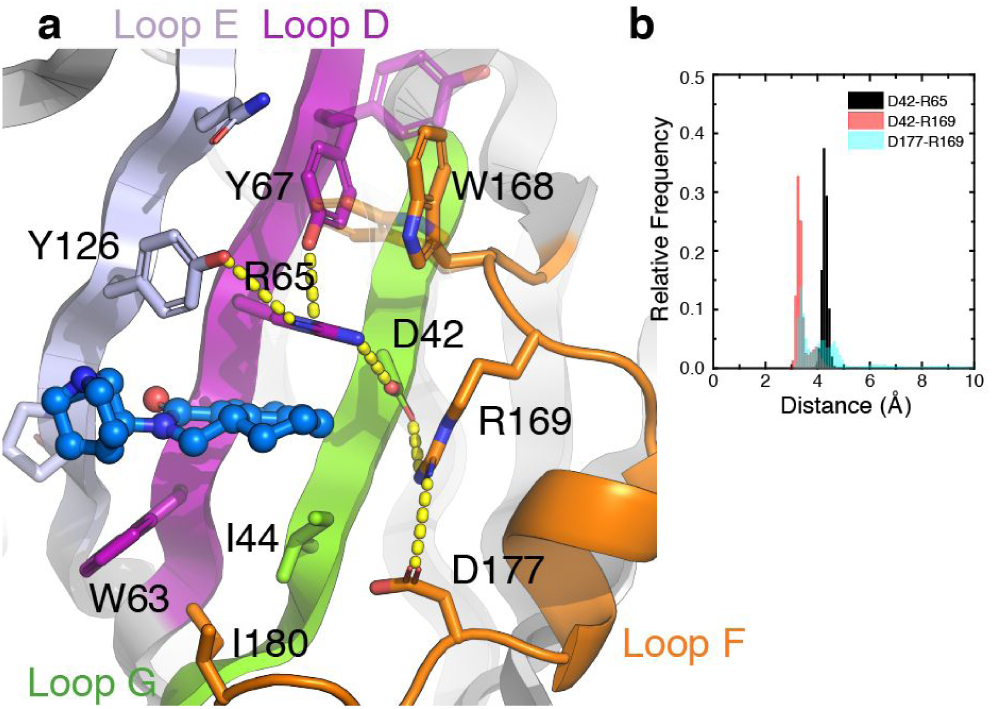
Network of side chain interactions on the complementary subunit. **a**. Cartoon and stick representation, approximately parallel to the membrane. Side chains within 4 Å of the drug are depicted as sticks. Yellow dashes indicate H-bonds or salt bridges. The dual conformation of residues Y67 and W168 (see Methods) is depicted as transparent sticks. **b**. Distance distributions between atoms forming the three salt bridges D42-R65, D42-R169 and D177-R169 (black, red and transparent cyan respectively) sampled during the MD simulation in the presence of palonosetron.

### Conformational dynamics of the drugs, H-bonding patterns

The protonated nitrogen of the azabicyclo ring forms a H-bond with the main chain carbonyl of W156 (Figure 7). This H-bond is consistently observed throughout the MD trajectory for four palonosetron molecules, whereas the fifth one engages in a long-lived H-bond with N101 (Figure 4b,c). The equivalent nitrogen of granisetron steadily interacts with W156 in four subunits and is left unpaired in the last subunit (Figure SI7). In trajectories generated for ondansetron and cilansetron, the nitrogen also forms H-Bonds with W156 and N101, with a preference for the latter residue (Figure SI9). This trend is more pronounced for ondansetron, whereas cilansetron also occasionally forms an H-Bond with S155 (Figure SI8). Dolasetron engages in a long-lived H-Bond with N101 through its ring nitrogen. It is also transiently H-Bonded to W156 but through its carboxyl oxygen, at variance with the other -setrons. Interestingly, dolasetron also forms an H-Bond between the nitrogen of its aromatic moiety and W63 (Figure SI6).

**Figure 4.**
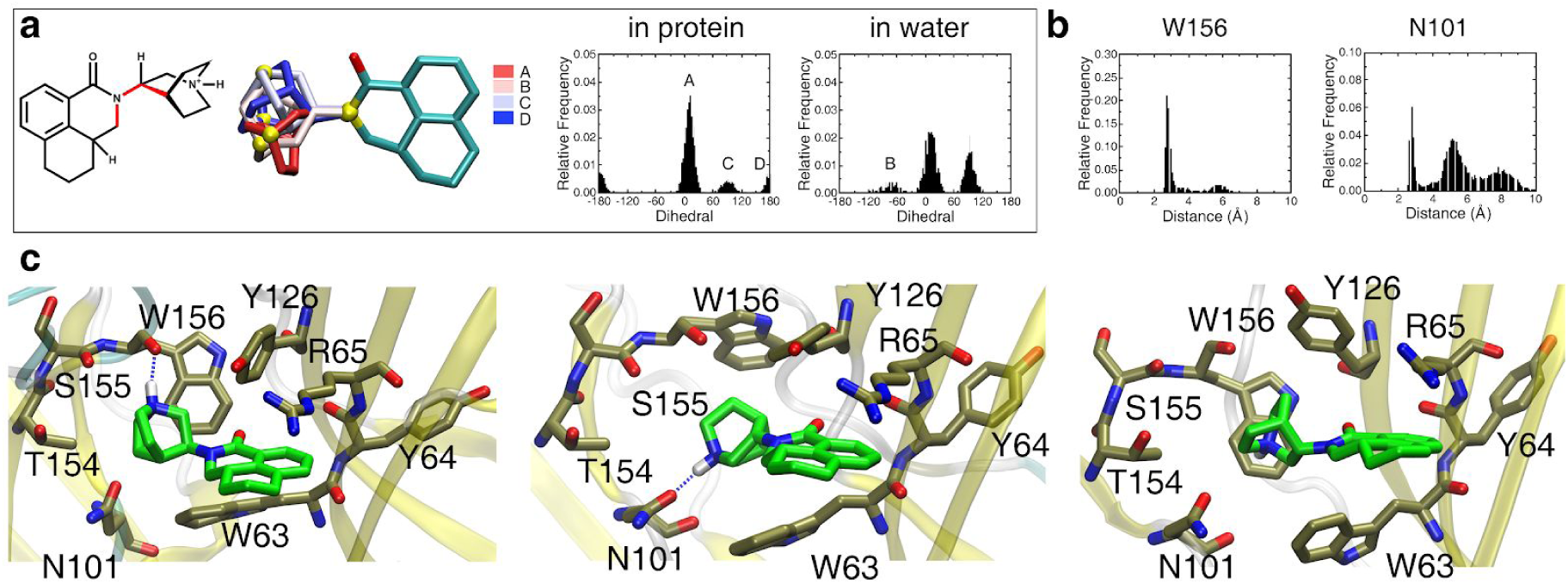
H-bonding pattern and dynamics of palonosetron during MD simulations. **a**. The inset shows the distributions of the dihedral angle between the two moieties of palonosetron (drawn in red), either bound to the receptor or free in water. The position of the aromatic part is relatively conserved in the protein indicating that the conformations sampled result from the conformational dynamics of the azabicyclo ring in the binding site. **b**. The two distance distributions depict the H-Bonds between the azabicyclo ring -NH group and the backbone carbonyl of W156 (the major interactant) and the sidechain carbonyl of N101 (note the sharp peak at 2.5 Å distance) monitored over the MD trajectory for the 5 subunits. **c**. Three snapshots illustrate the main conformations sampled by palonosetron.

The H-bond dynamics observed at the binding sites along the MD trajectories seem correlated with the conformational dynamics of the inhibitors (Figure 4a and SI6-SI9). It essentially involves a rotation around one (two for cilansetron) dihedral angle connecting the two rigid moieties of the -setron drugs, resulting in distinct conformations, with populations possibly modulated by the environment, which are observed for all drugs in solution or bound to the 5-HT3 receptor. The free-energy profile along the dihedral connecting the palonosetron rigid moieties, computed both in water and in the protein, revealed four minima (labeled A-D in Figure 4a) separated by barriers amounting up to 4 kcal/mol (Figure SI10). It echoes the 1993 report on the discovery of palonosetron, where the authors noted: *“An affinity of this magnitude could only be associated with a compound that binds at or near its global minimum”*^*19*^.

### What interacts with the ligand H-bond acceptor?

The presence of an H-bond acceptor in between the aromatic moiety and the cycle-embedded nitrogen moiety seems a defining feature of 5-HT3 receptor antagonists^20^. Yet, no obvious interaction for this chemical group is immediately visible when observing the palonosetron-bound cryo-EM structure (or the granisetron-bound one). In all simulations, we observed the formation of an H-bond between a water molecule and the oxygen of the ligand central carbonyl. This water molecule also engages in two H-bonds with the protein, one with the backbone carbonyl of Y64 and one with the -NH group of the W156 indole. In experimental densities, a faint peak is recorded and may correspond to a water molecule, this signal being stronger in the granisetron structure (visual inspection of the 6NP0 density map). We placed a water molecule and conducted new MD simulations. Indeed, this molecule remained trapped most of the time in the case of palonosetron (Figure 5), and at various levels for the other drugs (Figure SI11).

**Figure 5.**
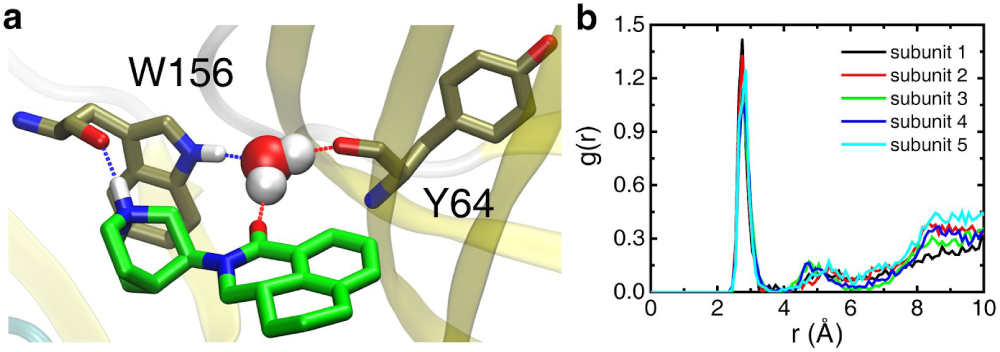
A trapped water molecule links palonosetron H-bond acceptor and the receptor. **a**. Representative snapshot illustrating the water mediated H-Bond network linking the palonosetron central carbonyl to protein residues W156 and Y64. **b**. Pair radial distribution functions between the oxygen of the palonosetron central carbonyl and the water oxygen, characterizing the probability density to find the two species at a given distance. The peak located at ∼3 Å indicates that for every subunit, there is always one water molecule H-bonded to palonosetron.

### A structural framework for functional data

Over nearly three decades, numerous studies have probed how mutations in, or around, the orthosteric binding site influenced anti-emetics inhibition of the 5-HT3 receptor. We briefly discuss the literature, in light of our structure and simulations, for each stretch of the receptor participating in the binding site.

#### Loop A

Among loop A residues, only N101 lies close enough to palonosetron (and to other ligands in other structures) for a direct interaction (Figure 6). In our simulations, the distance distributions of N101 with -setron drugs feature a main peak at 5 Å with two additional peaks at 2.5 Å and at ∼8 Å, indicating transient H-bonding of the ligands with this residue (Figures SI6-SI9). Palonosetron is 10-fold more potent with the N101A mutant, while N101Q remains similar to the wild type^11^, in line with the idea that the absence of H-bonding capacity would stabilize the azabicyclo moiety in the conformation where the N interacts with W156 only. Interestingly, mutations of N101 into a variety of amino acids (A, D, E, L, Q, R, V) does not affect [3H]-granisetron binding^21^. Accordingly, simulations show an absence of H-bond with the drug at this residue.

**Figure 6.**
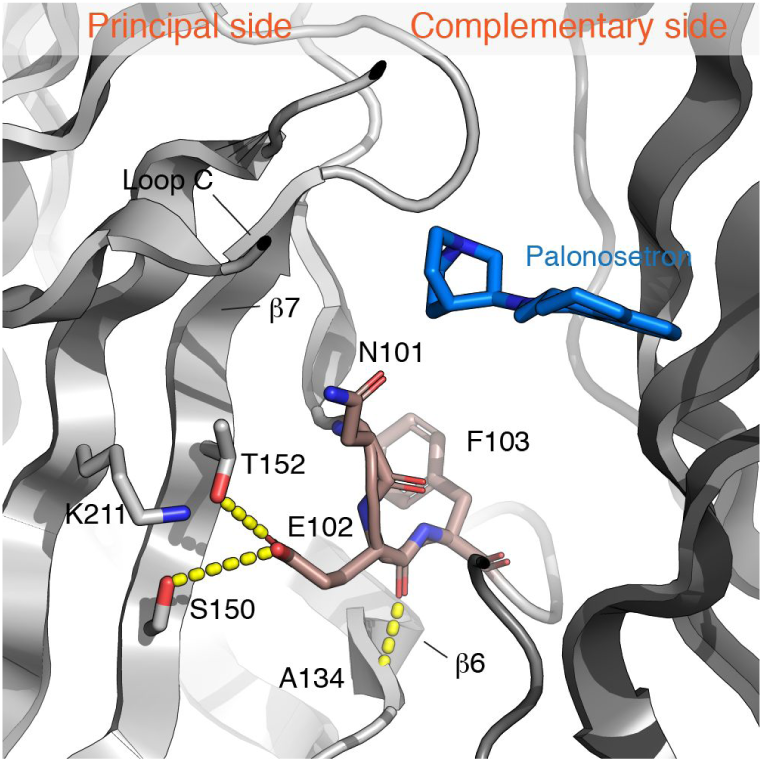
Loop A. On a cartoon representation, the three most important residues of loop A are shown as sticks, together with neighbors of E102. The interactions of E102 described in the text are represented as dashes.

**Figure 7.**
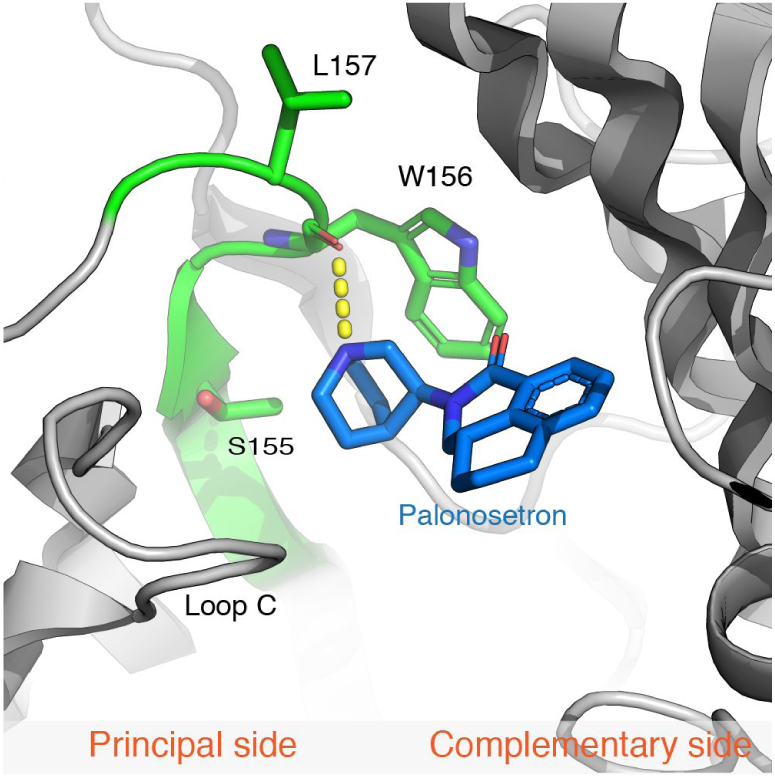
Loop B. On a cartoon representation, residues of loop B facing palonosetron (blue sticks) are shown as green sticks. The interaction of W156 main chain with the protonated nitrogen of the ligand is depicted as a dashed line.

[3H]-granisetron-binding experiments illuminate a drastic effect of mutations of E102, with a complete loss of binding at the subnanomolar range for A, D, G, K, N (data obtained on oocytes)^22^; or 60- to 80-fold decrease in K_d_ for E102D/N (data obtained on HEK cells)^23^. E102 is not oriented towards the binding site and interacts with S150 and T152 (β7), while its main chain carboxyl forms an H-bond with A134 (β6) (Figure 6). Thus, one possible role for E102 is to pack the β-sandwich and to position properly loops A and B. In addition, E102 creates an electronegative patch next to the binding site entrance, which might participate in ligand attraction, in line with the observation that the deleterious effect of introducing a net positive charge in loop A (e.g. N101K, E102Q) can be assuaged by removal of the positive charge at K211(regardless of H-bonding ability at position 211)^24^. At position 103, granisetron binding appears only marginally affected by F103Y/N substitutions^25^, while it was decreased ∼5-fold by F103A/W mutations^21^.

#### Loop B

Loop B harbors in most pLGIC a pivotal tryptophan residue (here W156), which forms the back wall of the binding site (Figure 7) and establishes cation-π interactions with ligands^26^. Unsurprisingly, alanine substitution abolishes granisetron and palonosetron binding^18,27^, while a more conservative mutation to tyrosine increases 10- and 180-fold the respective K_d_ of these drugs. Nearby alanine substitutions are relatively innocuous. L157A has a 10-fold effect on granisetron binding, maybe because the smaller side chain does not hinders the possible motions of Y126. L157I is wild-type-like.

#### Loop C

Loop C appears to be the most variable loop (with loop F) in multiple sequence alignments of pentameric ligand-gated channels. It undergoes deformation upon ligand binding, which remains modest in the 5-HT3 receptor structures. As a heritage from studies of stiff acetylcholine binding proteins (AChBP), where loop C was the most evident moving element^28,29^, its position is often referred to as a hallmark of the receptor state. Legitimate emphasis on loop C should not blunt other relevant markers of states (see below). Yet, it remains worth underscoring the simultaneous motion of the flexible loop C and ECD/TMD interface loops. Loop C connects by the rigid β9 and β10 strands, stabilized further by the N-linked carbohydrates (Figure SI12). The coupling between ligand binding and pore opening is a complex process^30,31^, but a structurally intuitive hypothesis is that motion of loop C in response to ligand binding might be transmitted to the ECD/TMD interface by means of a central β-sheet rigid hinge.

I201 and Y207 are the loop C residues that point towards palonosetron (Figure 8). Only aromatic mutations are tolerated at Y207, with Y207F yielding a 3-fold increase of K_d_ ^18^ (11-fold increase of the granisetron K_d_ ^32^). Interestingly an extra density is present at the tip of Y207 (at low contour level), which might correspond to a water molecule linking this residue to the complementary side via Y126 and Q124. I201 is not conserved in the human receptor. It corresponds to M228, the mutation to alanine of which produces a 6-fold decrease of K_d_ ^18^, in line with the van de Waals interaction observed in the structure (I201A also deteriorates 5-fold granisetron binding in the mouse receptor^32^). From a structural standpoint, loop C is stabilized by interactions of E198 and S206, and the presence of a negative charge at its tip (D202) might participate in attracting the azabicyclo moiety during binding. In line with a much broader spatial rearrangement of loop C following serotonin binding, mutations of loop C residues tend to have stronger effects on serotonin binding and activation than on binding of granisetron or palonosetron (for instance F199Y does not change the binding of both antagonists, but decreases the serotonin K_i_ by a 180-fold, while D202A entails a 140 K_i_ decrease and 12-fold EC_50_ increase).

**Figure 8.**
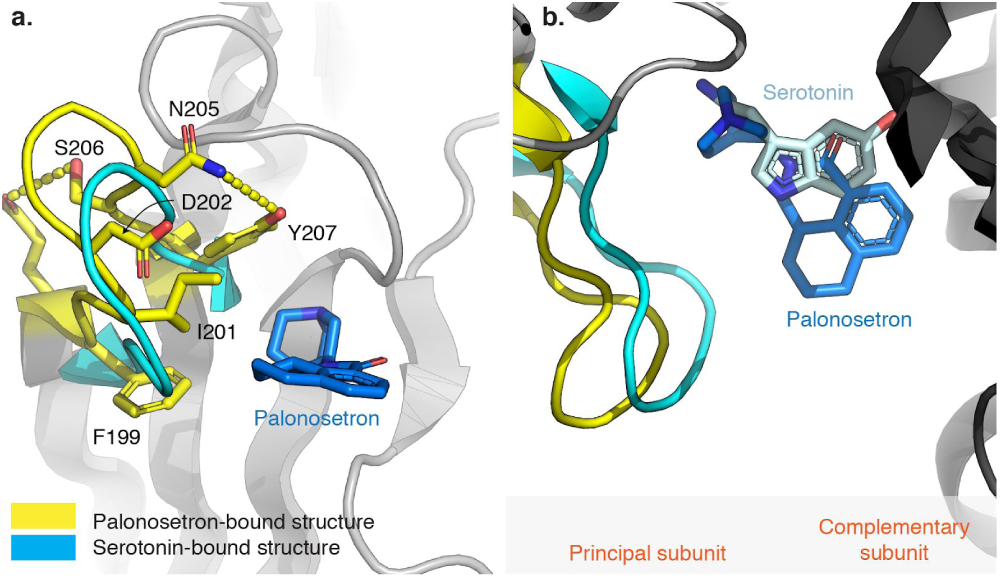
Loop C. The serotonin-bound activated structure is superimposed to the palonosetron inhibited one (on the complementary subunit) to highlight the yellow→cyan capping motion of loop C. **a**. View parallel to the membrane plane of loop C. Palonosetron and residues which mutations are discussed in the text are represented as sticks **b**. View perpendicular to the membrane plane. Here, serotonin present in the activated structure is also represented as sticks.

#### Loop D (and G)

Two residues of loop D, W63 and R65, are in direct contact with palonosetron, sandwiching its isoquinoline ring (Figure 3). As mentioned previously, W63A is nonfunctional, while W63Y exhibits wild-type properties both for serotonin activation and palonosetron inhibition^11^. The conformation of R65 seems in line with a cation-π interaction both in our palonosetron-bound structure and in the granisetron-bound one^16^, and this interaction has been demonstrated experimentally with fluorinated derivatives of granisetron^10^. It is noteworthy that both R65A and R65C mutations yield different effects for different antiemetics. R65A reduces the binding affinity for granisetron, but produces no significant change for palonosetron^11,33^. R65C abolishes granisetron binding, yet marginally affects binding of tropisetron^34^. The differential effect was linked to putatively different orientations of the aromatic moieties of the drug, which is not the case for palonosetron and granisetron. In our previous, lower-resolution structure in complex with tropisetron^13^, we modeled its aromatic ring perpendicular to the ones of palonosetron or granisetron. However, ligand building in 4.5-Å maps is tricky, and better data would be required to ascertain this difference in orientation.

In loop G, D42C abolishes tropisetron binding, albeit barely modifies that of granisetron^34^.

#### Loop E

Mutations at Y126 affect both granisetron (Y126A, 8-fold)^35^ and palonosetron binding (Y126A/F, 20-30-fold)^18^, in line with the proximity of the side chain to palonosetron and the implication of the terminal hydroxyl in intra and inter-subunit interactions (Figure 3). The necessity of H-bonding capacity at Y126 has also been established by nonnatural amino-acid mutagenesis^36^.

In addition, mutagenesis suggests roles of neighboring residues in shaping the binding site properly, as alanine mutation of any residues in the 124-127 range produces a 6 to 8-fold decrease in the granisetron K_d_ ^35^.

#### Loop F

The extensive network of interactions on the complementary side, together with mutagenesis, points to an essential role of the precise positioning of loop F. Mutants at positions W168(Y), P171(H), L167(A), W168(A), R169(A) and K178(A) yield >5-fold increase of granisetron K_d_, while D177(A) shows no affinity^37^ (Figure 9). The interaction of D42, R169 and D177 readily explains the mutant phenotypes at these positions: if disrupted, free charged side chains would impede the drug binding. Interestingly enough, D177N/E mutations, preserving the H-bonding capacity, yield unchanged granisetron binding, while D177A abolishes it. Serotonin activation is also lost for D177A. On top of its interaction with R169, D177 also positions the bottom of loop F through an interaction with the main chain of S179. This idea that the precise position of loop F is crucial for function is further reinforced by the effect at K178, which anchors the loop to the β9 strand at the main chain of L190. Conversely, mutation of R175 does not produce any change, in line with its orientation towards the solvent.

**Figure 9.**
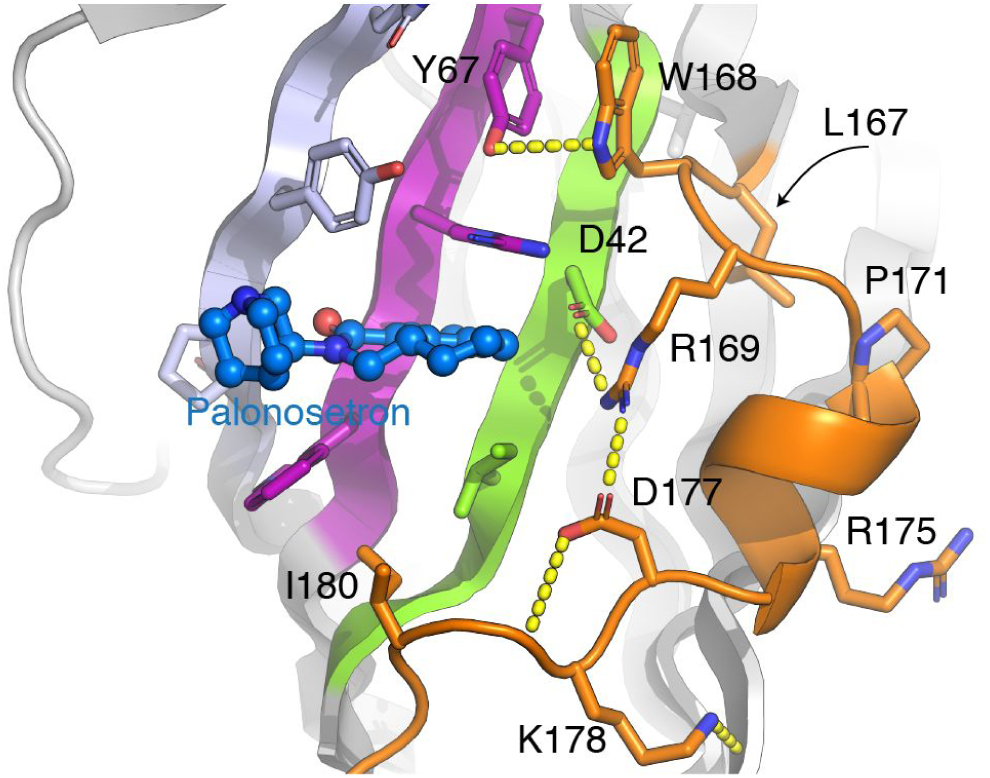
Loop F. On a cartoon representations, loop F residues (orange) which mutations are discussed in the text are represented as sticks.

### Would palonosetron bind at heteromeric interfaces?

Five homologous subunits of the 5-HT3 receptor are encoded by the human genome, while rodents possess only two of them. In heteromeric receptors, the activation interface is usually thought to be only the A(+)/A(-) one, i.e., the one where palonosetron fits tightly in the present structure. Early studies have concluded that some ligands bind differently to different subpopulations of receptors^38^, and one wonders whether palonosetron may also bind to other interfaces. Mutations at position W156, R65 and Y126 yield no or only poor binding of palonosetron. Thus, one can infer that interfaces involving a B(+), C(+), D(+) or E(+) subunit would not bind the drug (they all lack the tryptophan in loop B, at the position equivalent to W156). Similarly B(-), C(-) or E(-) interfaces lack an arginine at positions equivalent to R65, and D(-) interfaces lack a tyrosine at position equivalent to Y126. Put together, we hypothesize that only A(+)/A(-) interfaces are responsible for palonosetron binding and inhibition.

### The singularity of palonosetron?

Discovered in 1993^19^ and marketed in 2003, palonosetron was coined a “second-generation antagonist”, obviously because it came after granisetron, tropisetron and ondansetron, but also because, unlike these molecules, it was reported to prevent not only acute-, but also delayed-CINV in clinical studies^39^. Palonosetron distinctive features (higher potency^40^ and longer plasma half-life^41^) cannot in principle explain this unique effect on delayed CINV: other drugs administered more often and at higher doses would also prevent it. In attempts to identify other singularities that might rationalize the advertised clinical outcome, it was also proposed that palonosetron 1) shows cooperativity and binds to allosteric sites^42^, 2) triggers 5-HT3 receptor internalization^43^ and 3) influences the crosstalk between 5-HT3 receptors and the neurokinin 1 receptor in a way that elicits a synergistic effect of antagonists at each receptor^44^ (neurokinin 1 inhibitors such as aprepitant are widely used in combination with -setron drugs for CINV).

Our data unequivocally shows that palonosetron binds to the orthosteric site; no secondary site was apparent from the inspection of the residual densities, in line with mutagenesis experiments disproving the existence of an allosteric site underneath the neurotransmitter one^18^. Granisetron, tropisetron, and palonosetron bind to the same site and apparently stabilize the same conformation of the homomeric 5-HT3A receptor: In this respect, the clinical singularity of palonosetron is not apparent in current structural studies.

We have however underscored some of the palonosetron unique features in terms of binding and dynamics: the preeminence of its conformation of minimal energy (Figure 4a, SI10); the continuous presence of a water molecule next to its H-bond acceptor (Figure 5). In addition, the notion of *“restricted”* conformational freedom of the aromatic moiety, which guided the synthesis of palonosetron^19^, is clearly echoed here. The distribution of the angle between the aromatic moiety and the main axis of the protein is narrower for palonosetron compared to other drugs (Figure SI13).

### Serotonin-bound ECD conformations clearly differ from antagonist-bound ones

The orthosteric site of pentameric ligand-gated channels lies at the interface between subunits, and quaternary reorganization of extracellular domains is key to their activation^13^. To get a sense of the spread of quaternary arrangements of available 5-HT3 receptor structures, we superimposed their complementary subunits and looked at their principal subunits (Figure 10). Two groups are clearly distinguished. The average pairwise RMSD within the group of inhibited structures is 0.49±0.06 Å, while the average pairwise RMSD between inhibited and activated structures is 0.91±0.22 Å. At the binding site level, the activated structures possess tighter cavities where -setron binding would not be possible (Figure 10c).

**Figure 10.**
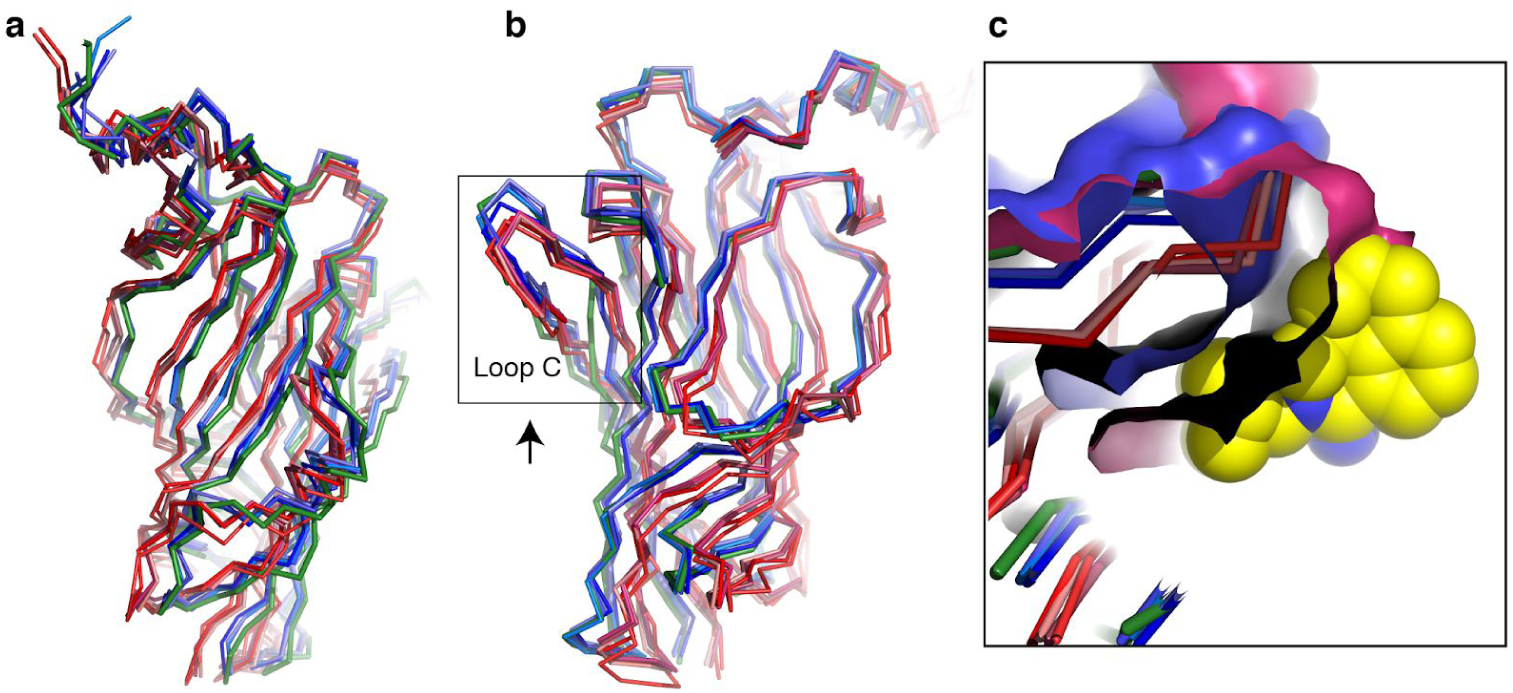
The inhibited conformations form a group that differ from the active ones. Because the quaternary organization is key to the gating of the 5-HT3 receptor, we here represent the principal subunit ECD after alignment of the complementary subunit ECD. Active conformations are depicted in red, inhibited in blue, while the apo one is in green. All the inhibited conformations are clearly grouped, all the active ones as well. The inhibited group is closer to the apo conformation, and similar at the ECD/TMD interface while differences are seen in β8 and β9. **a**. and **b**. panels are two different orientations where the subunit is seen from the point of view of its two neighbors. **c**. Palonosetron (yellow spheres) fits tightly in its binding site (surface of loop C in blue) but would clash with the binding site of the active conformation (red surface of loop C).

The apo structure rather lies within the cluster of inhibited ones, following the allosteric theory hypothesis stating that ligands stabilize preexisting conformations^45^. However, as already noted by Basak et al.^16^, the apo structure presents moderate deviations compared to the inhibited ones, which are most apparent at the level of strands β8 and β9, and of loop F. In line with the closed phenotype in the absence of agonist, no deviation is observed at the ECD/TMD interface, a key region for the transduction of the signal from the orthosteric site to the pore gate.

### 5-HT3 receptor inhibitors beyond emesis?

5-HT3 receptor antagonists are widely used for the control of emesis. Their potential in other therapeutic areas, by themselves or as adjunct drugs, has been explored in preclinical and clinical studies with equivocal results. A significant fraction of these studies were carried out more than two decades ago and it is worth combining older^46^ and more recent reviews^4,5,47^ to get a sense of the yet unmet promises of 5-HT3 receptor inhibition in areas such as depression, anxiety, schizophrenia and drug withdrawal.

In parallel, roles of 5-HT3 receptors in a variety of tissues continue to be uncovered, for instance in the lacrimal gland^48^, in the development of the urinary system^49^, as well as in adipocytes^7^. These and future similar studies may one day help reinterpret older observations from (pre)clinical studies. In our era of drug repurposing, -setron molecules, which enjoy a good safety profile and a relative absence of adverse effects, might spur renewed interest.

### Conclusion

Detailed knowledge of antiemetics mode of action with the 5-HT3 receptor has remained hitherto fragmentary. We have solved using Cryo-EM the palonosetron-bound structure of a mouse 5-HT3 receptor, which amid the very few related structures available, provides valuable new information towards understanding inhibitory mechanisms at the atomic level. Combining our structural and computational data, we put forth a convenient framework for rationalizing the host of biochemical and pharmacological information accrued since the first-setron drug synthesis. This framework is well-suited for correlating specific mutations to observed phenotypes, and for establishing a general structure-activity relationship for antiemetic drugs. As more ligand-bound structures of serotonin 5-HT3 receptors will likely be obtained at increasing resolutions, our grasp of pharmacology and conformational equilibrium will deepen. In the coming years we also anticipate that experimental data on asymmetry (either in homomeric or heteromeric receptors) and on conformational variability will continue to reshape the current view on the modus operandi of these fascinating allosteric neuroreceptors.

## Methods

No statistical methods were used to predetermine sample size. The experiments were not randomized. The investigators were not blinded to allocation during experiments and outcome assessment.

### Protein production and purification

The wild-type mouse 5-HT3A receptor was expressed in a stable and inducible cell line, and purified in the C12E9 detergent as previously described^13,50^.

### Electron microscopy and image analysis

The most homogeneous fractions after size-exclusion chromatography were concentrated to ∼1.5 mg/ml, mixed with lipids (0.01% phosphatidic acid, 0.01% cholesterol hemisuccinate, 0.01% brain phosphatidylcholine; Avanti Polar Lipids) and 1 mM palonosetron (gift from C. Ulens) and incubated for 10 minutes on ice. 3.5 μl were deposited on a glow-discharged (25 mA, 50 s) Quantifoil copper-rhodium R 1.2/1.3 grid, blotted at 8°C for 10 s at force 0 using a Mark IV Vitrobot and plunge-frozen in liquid ethane. The dataset was recorded on a Titan Krios electron microscope at the ESRF^51^(Table SI2). Movies (40 frames) were acquired with EPU on a Gatan K2 Summit camera mounted after an energy filter (20eV slit width).

The 3’037 raw movies were aligned and summed with MotionCor2^52^, within Relion 3.08^53^. CTF estimation for non dose-weighted sums was calculated using GCTF^54^. Particles were picked with the FPM algorithm^55^ using as templates 2D projections of the serotonin-bound 5-HT3 receptor (PDB code: 6HIQ), low-pass filtered at 30 Å. All subsequent steps were performed in Relion. A total of 514’057 images were extracted, 4×4 binned and sorted in 20 classes by 2D classification. Class averages that resembled a pentameric ion channel were selected, amounting 348’500 particles. Selected particles were sorted by 3D classification without imposing symmetry and using as a reference model the low-pass filtered 6HIQ map. The classes that presented typical 5-HT3 receptor features and some density for the ICD were selected. The selected subset, amounting 198’076 images, was re-extracted at full size and used for 3D auto-refinement with imposed C5 symmetry, resulting in an initial map at 3.6 Å resolution. The particles from this refinement job were further sorted in four classes by focused 3D classification, with a mask around the ICD, without image alignment. The best class, amounting 149’317 particles, was subjected to cycles of 3D auto-refinement, Bayesian polishing and CTF refinement resulting in a 3.2 Å unmasked map.

The ‘shiny’ particle stack from the last polishing job was imported to, and final steps were performed in, cryoSPARC^56^. After 2D classification cleaning (Figure SI14), 127’971 particles were used for an *ab initio* reconstruction, followed by three rounds of Non-Uniform refinement, with C5 symmetry, yielding a final masked map at 2.82 Å resolution. Local resolutions were estimated with the ResMap software^57^ (Figure SI14).

### Simulation details

The cryo-EM structure was the initial point for the simulation of palonosetron. A pure palmitoyl-oleyl-phosphatidylcholine (POPC) bilayer, consisting of 120 units in each leaflet, was generated using the CHARMM-GUI interface^58,59^. The complex was solvated with 38386 TIP3P water molecules^60^ and 150 mM KCl. The initial structures of granisetron-, ondansetron-, dolasetron-, and cilansetron-bound 5-HT3 pentamers were obtained by replacing the palonosetron by these ligands. The aromatic rings of the -setron ligands were aligned with the aromatic plane of palonosetron. The parameters for all ligands were generated by CHARMM general force field (CGenFF)^61^. The protein and lipid were described by CHARMM36 protein ^62,63^ and lipid^64^ force fields, respectively. Hydrogen mass repartition ^65^ was used for all drug-receptor complex simulations, allowing for using a time step of 4 fs.

Program NAMD 2.13^66^ was employed for all the simulations. The temperature and pressure were maintained at 1 atm and 300K by the Langevin piston method^67^ and the Langevin thermostat, respectively. Long-range electrostatic interactions were evaluated by the particle-mesh Ewald (PME) algorithm^68^. Integration was performed with a time step of 8 and 4 fs for long- and short-range interactions, respectively, employing the r-RESPA multiple time-stepping algorithm^69^. The SHAKE/RATTLE^70,71^ was used to constrain covalent bonds involving hydrogen atoms to their experimental lengths, and the SETTLE algorithm^72^ was utilized for water. For each molecular assembly, a 100 ns pre-equilibration was performed using soft harmonic positional restraints on heavy atoms, followed by a 400 ns production run with no restraints. The VMD program was employed to analyze and render the MD data^73^.

### Model refinement and structure analysis

Cycles of real-space refinement in Phenix^74^ were performed, alternating with manual rebuilding in Coot^75^. The model comprises residues 8–329 and 397–459. Palonosetron (numbered 903) was placed in the orthosteric site. Even though uncommon at this resolution, residues 67 and 168 were modeled in two alternate conformations (see text and Figure SI4) with occupancies set at 0.5. The model is 5-fold symmetric, as the reconstruction used to build it possesses C5 symmetry. Validation was performed with Molprobity ^76^ (Table SI2). Visual information about the model and its fit in the map are found in figures 1a,b, 2b, SI1, SI4, SI12 and SI15. Figures were prepared with PyMOL (Schrodinger), Chimera^77^ or **<u>CueMol</u>**.

## Data availability

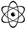 PDB 6Y1Z

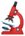EMD-10673

## Acknowledgements

We acknowledge access to ESRF CM01 Krios microscope. We thank L. Estrozi and A. Desfosses for advice in image analysis; members of the Nury group for discussions. The work was funded by ERC Starting grant 637733 Pentabrain. It used the platforms of the Grenoble Instruct-ERIC center (ISBG; UMS 3518 CNRS-CEA-UGA-EMBL) within the Grenoble Partnership for Structural Biology (PSB), supported by FRISBI

## Supplementary information

**Figure SI1.**
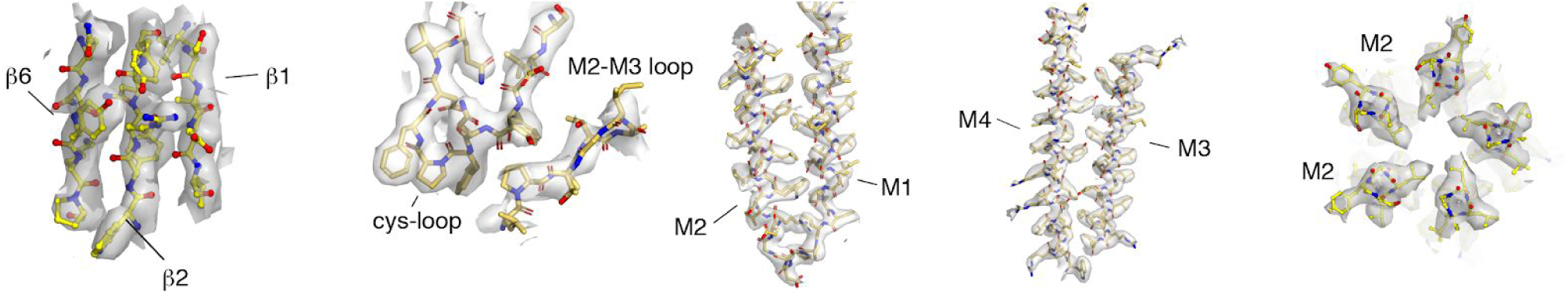
Quality of the density maps in representative parts of palonosetron/5-HT3 receptor reconstruction. Densities at 5σ in surface representations overlaid with the structure. Views a–e are approximately the same as those in previous publications13,78, to enable comparison. From left to right: densities of the β-sheets in the ECD, densities of the Cys loop and the M2–M3 loop, densities of helices M1 and M2, densities of M3 and M4, densities of M2 at the level of L9′ (L260).

**Table S1.**
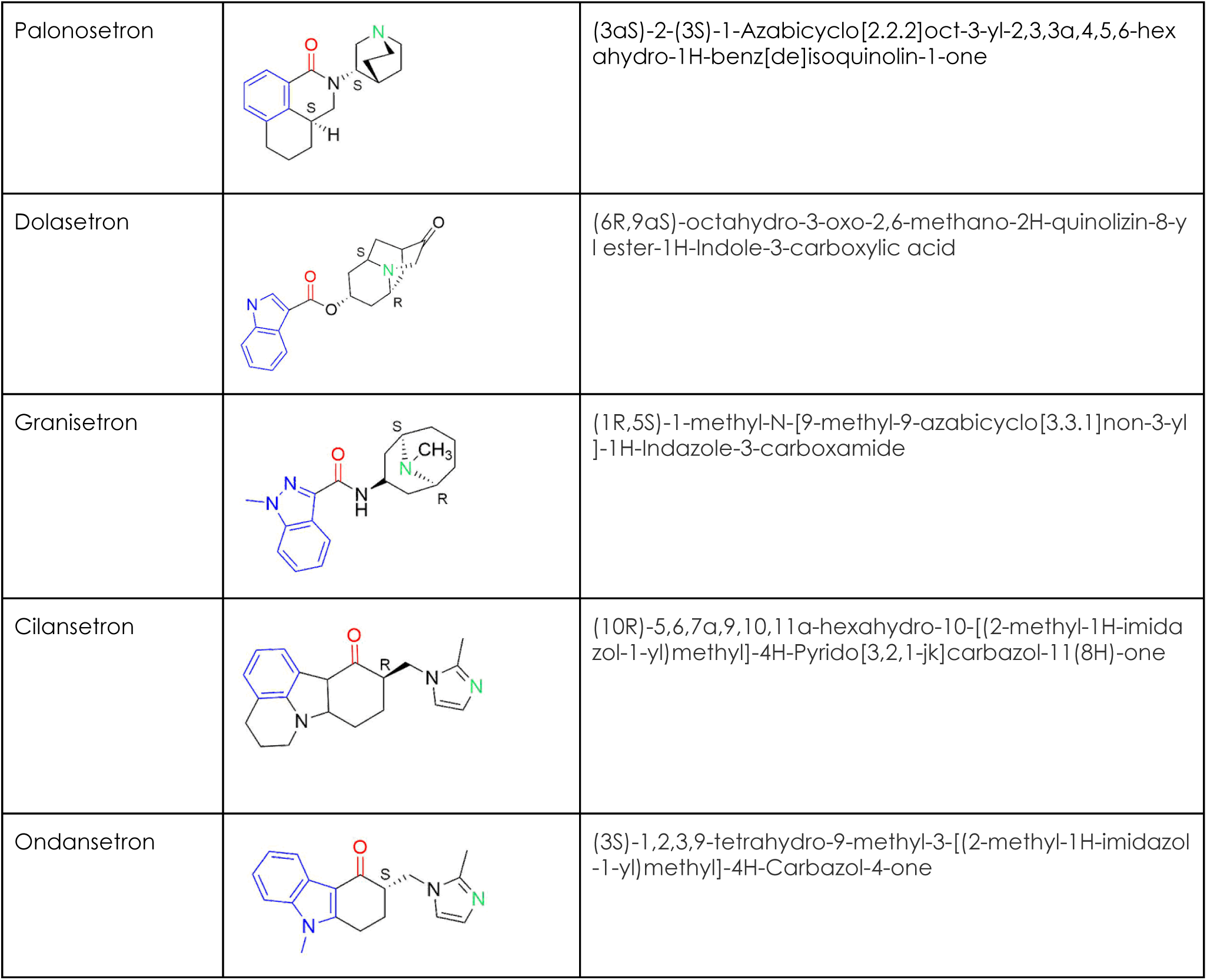
5-HT3 receptor antagonists used clinically. Each compound contains an aromatic moiety (blue), a ring-embedded protonated nitrogen (green) and an H-bond acceptor oxygen group (red). Isomers used in clinical formulations are indicated on the molecule schemes. The same isomers have been used in the MD simulations. For Ondansatron, the clinical formulation is a racemic mixture, only the S isomer is represented in the table.

|  |  |  |
| --- | --- | --- |
| Palonosetron |  | (3a*S*)-2-(3*S*)-1-Azabicyclo[2.2.2]oct-3-yl-2,3,3a,4,5,6-hexahydro-1H-benz[de]isoquinolin-1-one |
| Dolasetron |  | (6*R*,9a*S*)-octahydro-3-oxo-2,6-methano-2H-quinolizin-8-yl ester-1H-Indole-3-carboxylic acid |
| Granisetron |  | (1*R*,5*S*)-1-methyl-N-[9-methyl-9-azabicyclo[3.3.1]non-3-yl]-1H-Indazole-3-carboxamide |
| Cilansetron |  | (10*R*)-5,6,7a,9,10,11a-hexahydro-10-[(2-methyl-1H-imidazol-1-yl)methyl]-4H-Pyrido[3,2,1-jk]carbazol-11(8H)-one |
| Ondansetron |  | (3*S*)-1,2,3,9-tetrahydro-9-methyl-3-[(2-methyl-1H-imidazol-1-yl)methyl]-4H-Carbazol-4-one |

**Figure SI2.**
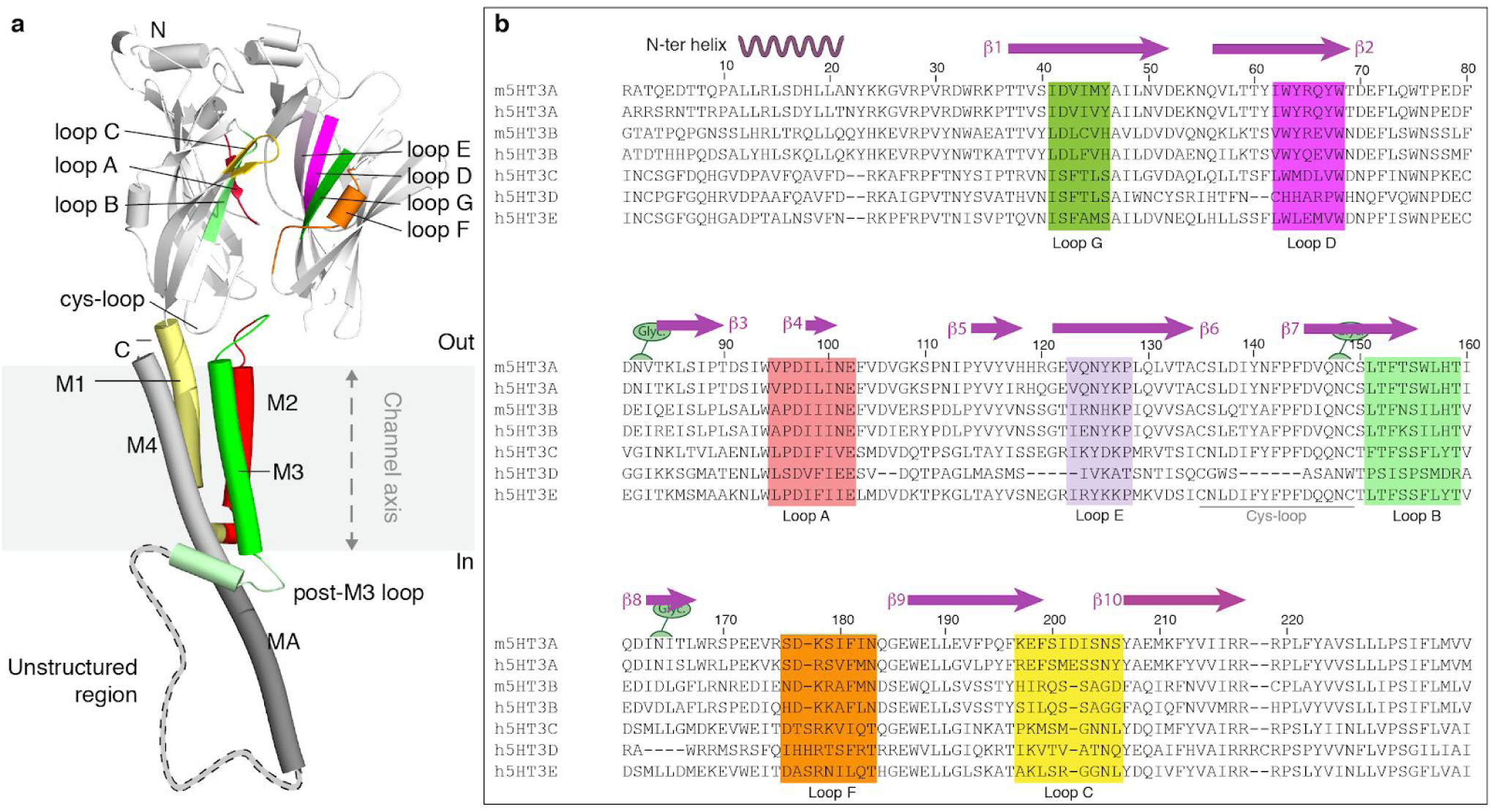
Nomenclature of the orthosteric site and sequence alignment. **a**. Ribbon representation of a full subunit, and two adjacent extracellular domains, with colored stretches for the binding elements making up the neurotransmitter site: 3 loops named A-C on the principal subunit, 3 strands and a loop named D-G on the complementary subunit. **b**. Multiple sequence alignment of the mouse and human 5-HT3 receptors, with binding elements highlighted.

**Figure SI3.**
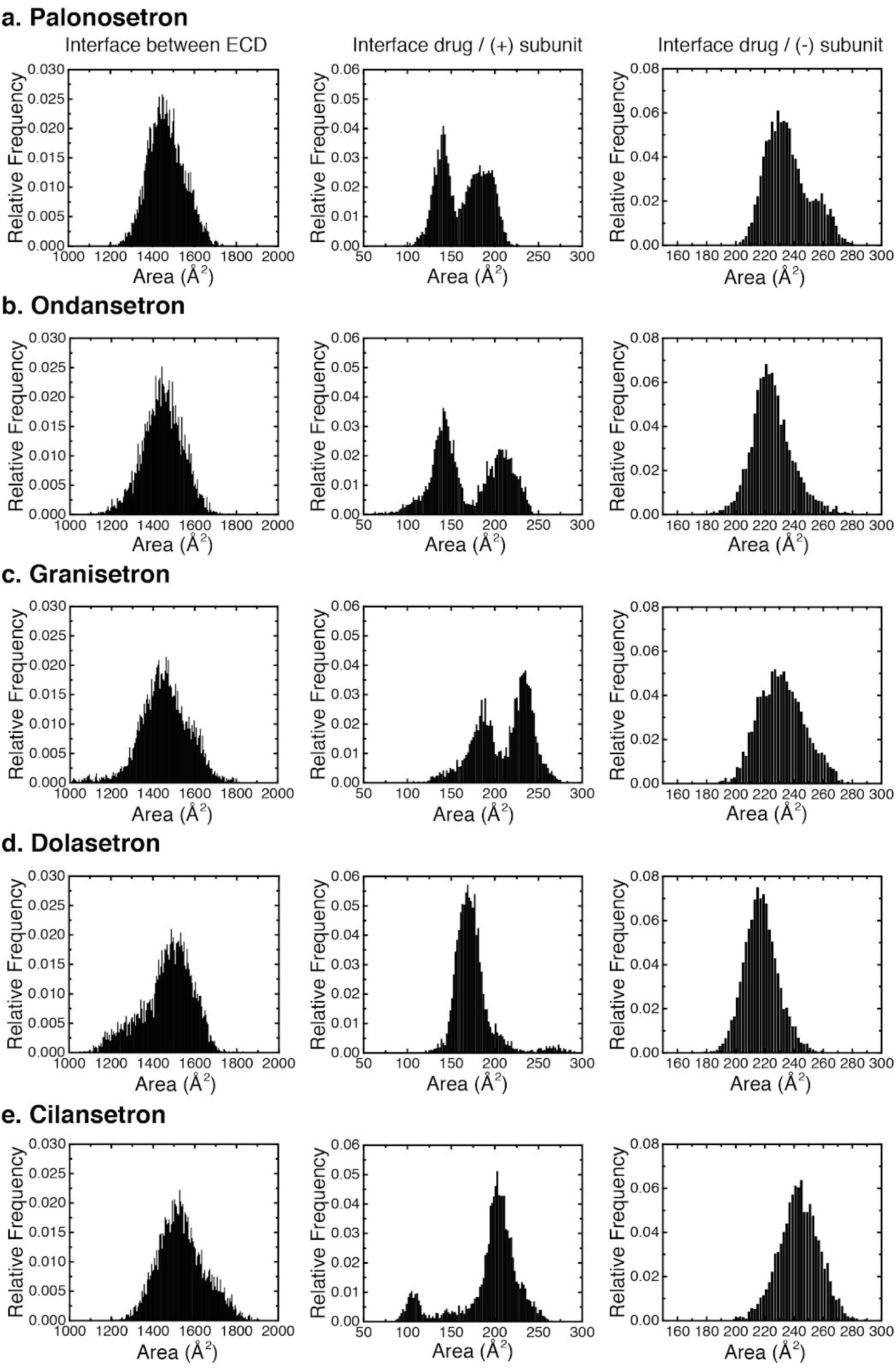
Normalized distribution of interaction areas sampled along the MD trajectories. Left column. Interface area between ECDs. **Middle column**. Interface area between the drug and the complementary subunit. **Right column**. Interface area between the drug and the principal subunit.

**Figure SI4.**
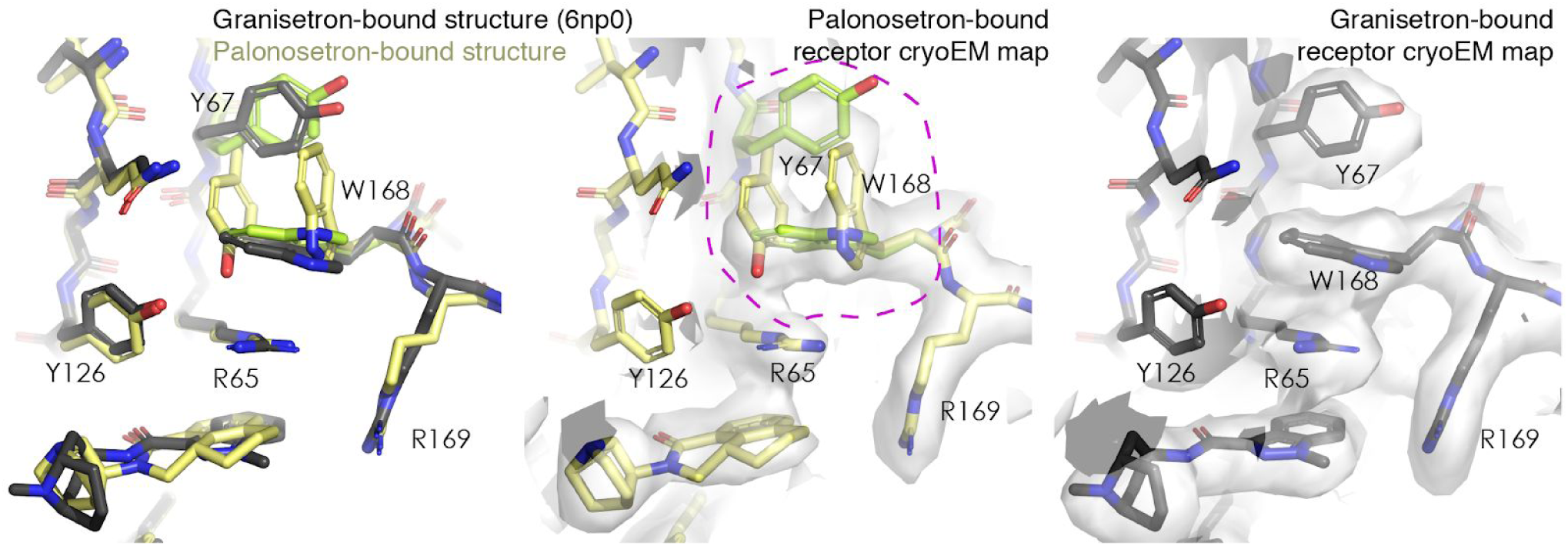
An alternate conformation at the level of Y67 and W168? Above R65, the density map appears more noisy than in the surroundings (middle panel, zone encircled in purple), which may indicate that W168 and Y67 and W168 can adopt dual conformations. The first one (represented in black from the 6np0 granisetron-bound model, or in limon) features a four-layer stack (drug-R65-W168-Y67) where Y67 lies far from the drug, while in the second one (represented in yellow), W168 moves out to permit a direct interaction of Y67 with R65. Mutagenesis seems to indicate that the ability to form hydrogen bond at position 67 impacts palonosetron binding (Y67F shows a 6-fold decrease in Ki)^11^. We ran molecular dynamics simulations of the two possible palonosetron-bound conformations and observed a better stability of the drug in conformation #1. In the granisetron-bound map of Basak et al. (depicted in the right panel), only the conformation #1 seems present. Cysteine substitution at position 67 affects slightly granisetron binding but drastically tropisetron binding^34^, implying that the relative position of loop D side chains and of the drug aromatic moiety varies from drug to drug.

**Figure SI5:**
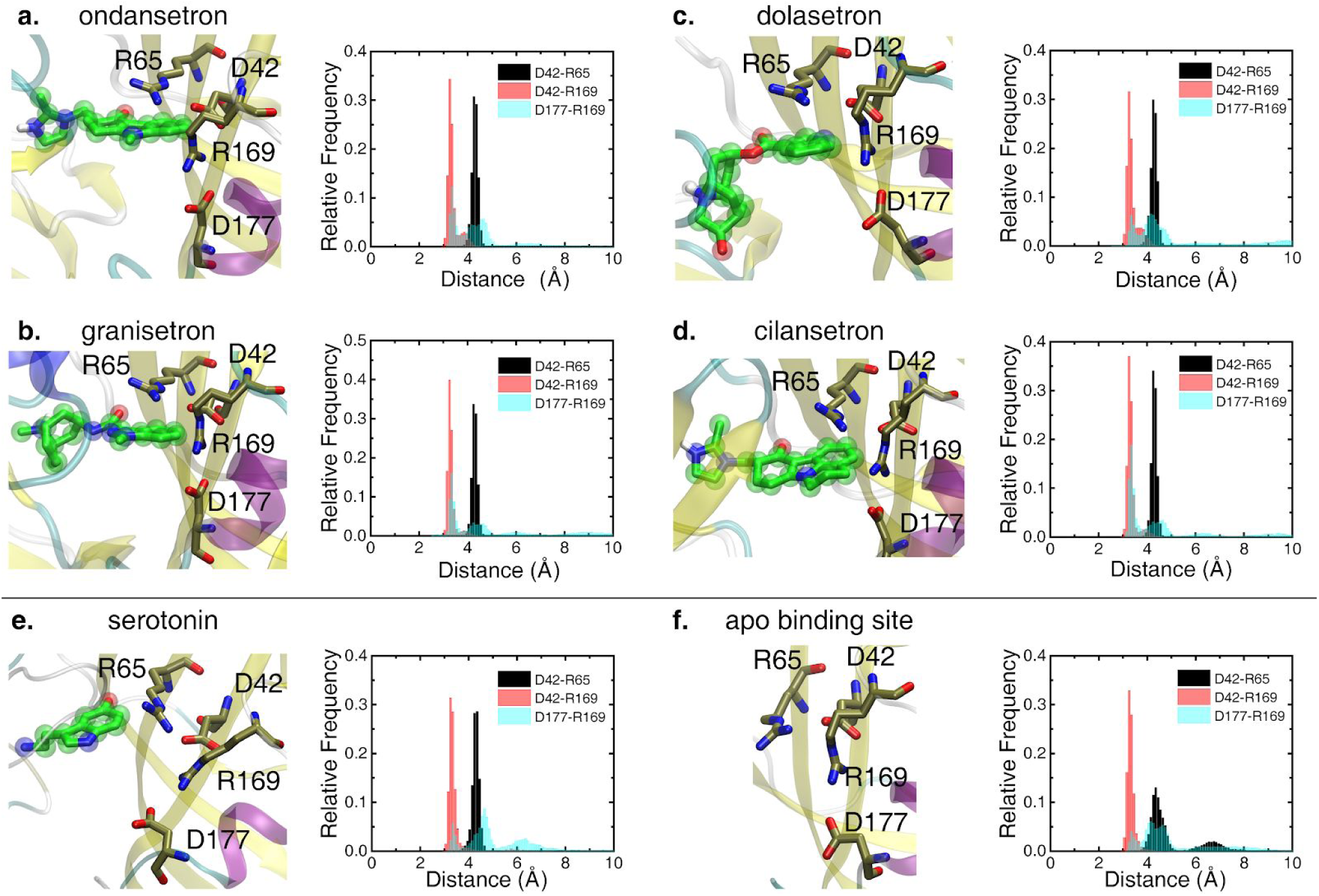
Evolution of the salt bridges during the MD trajectories. Representative snapshots and distance distributions between atoms forming the three salt bridges D42-R65, D42-R169 and D177-R169 (black, red and transparent cyan respectively) sampled during the MD simulation in the presence of (a) ondansetron, (b) granisetron, (c) dolasetron, (d) cilensentron, (e) serotonin and (f) without drug.

**Figure SI6.**
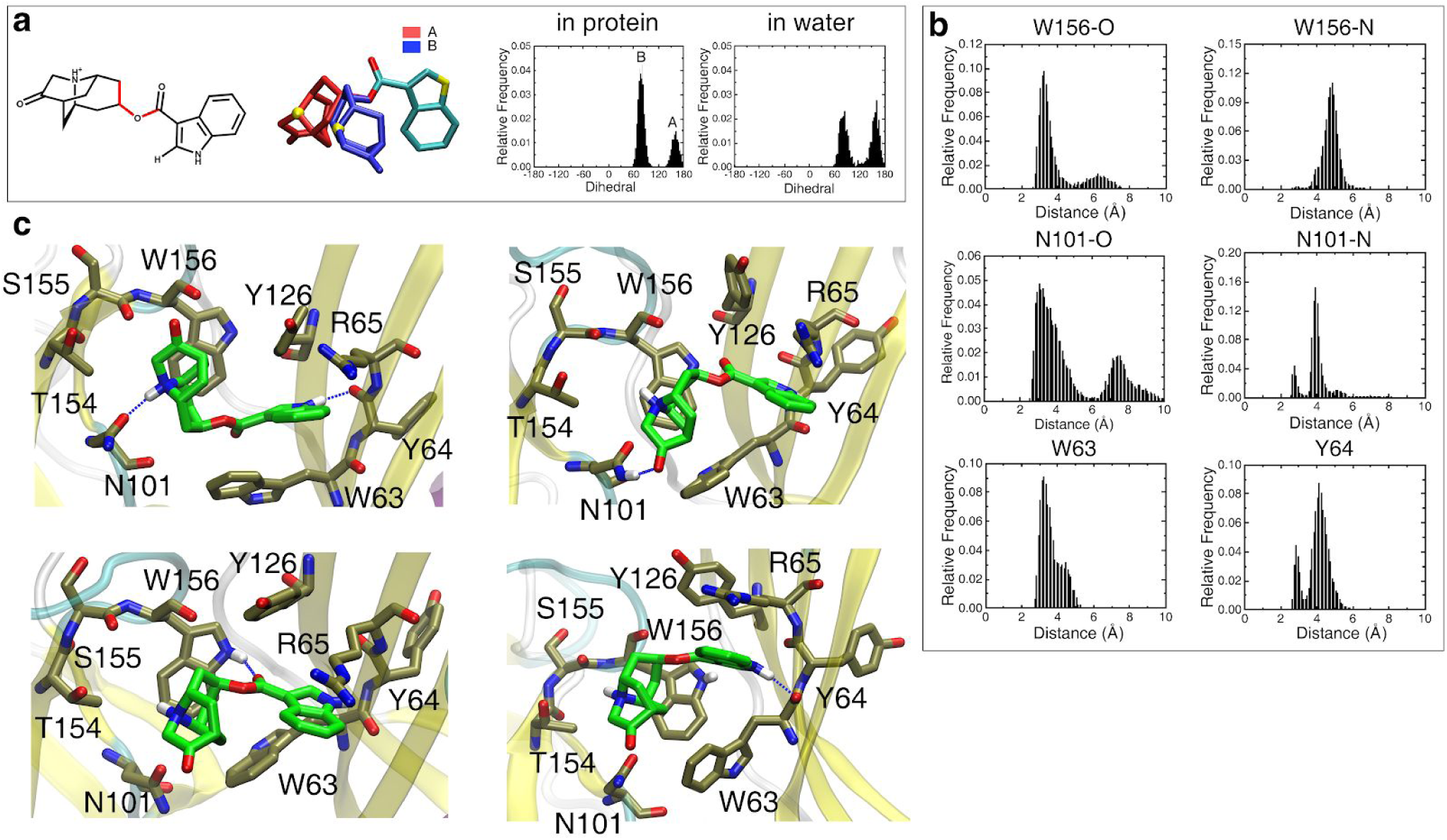
H-bonding pattern and dynamics of dolasetron during MD simulations. **a**. The inset shows the distributions of the dihedral angle between the two moieties of dolasetron (drawn in red), either bound to the receptor or free in water. The position of the aromatic part is relatively conserved in the protein indicating that the conformations sampled result from the conformational dynamics of the azabicyclo ring in the binding site. **b**. The six distance distributions depict the H-Bonds between the dolasetron central carbonyl and the -NH group of W156 (W156-O), the ring -NH group and the backbone carbonyl of W156 (W156-N) or the sidechain carbonyl of N101 (N101-N), the carbonyl of the nitrogen cage and the-NH group of N101 (N101-O) and the -NH group of the dolasetron aromatic ring and the carbonyl of W63 or Y64 monitored over the MD trajectory for the 5 subunits. **c**. Four snapshots illustrate the main conformations sampled by dolasetron.

**Figure SI7.**
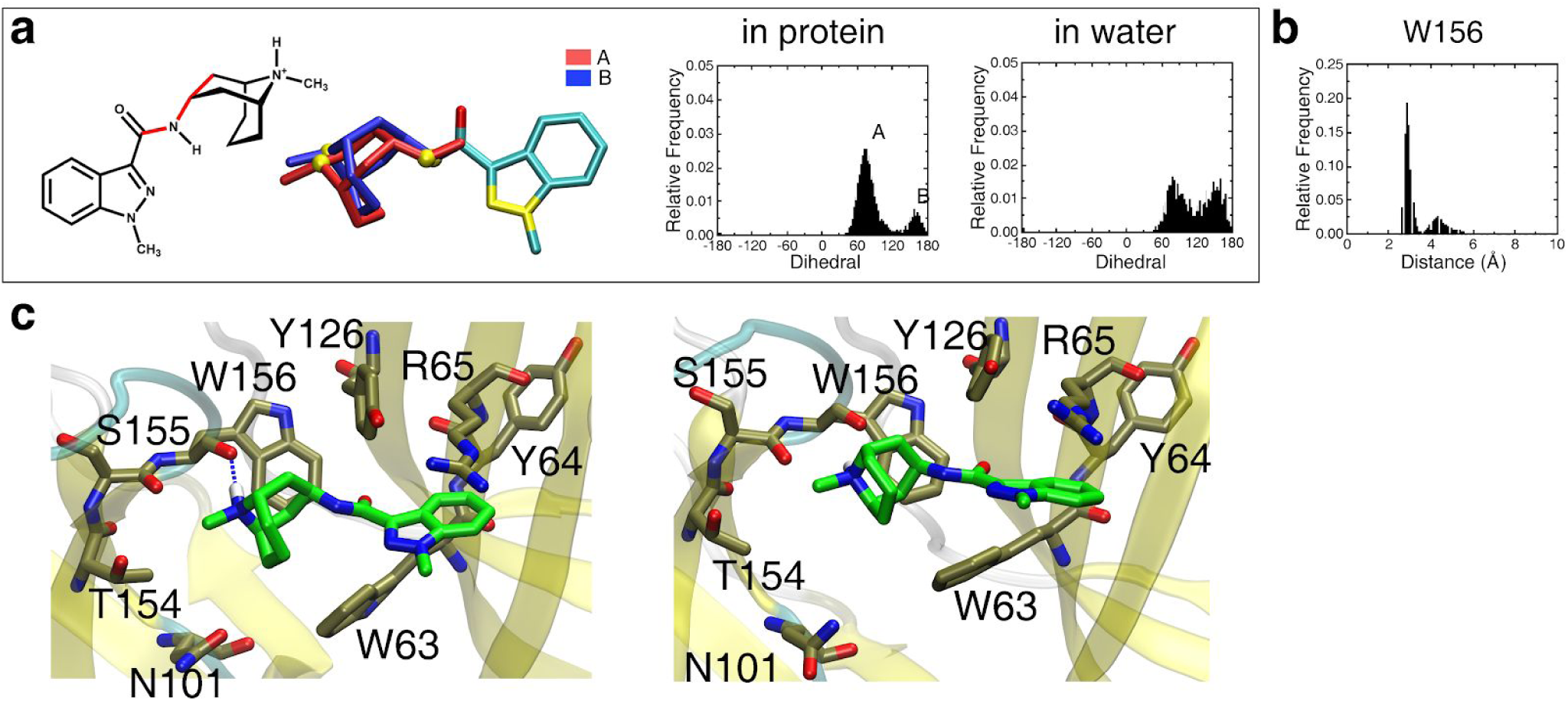
H-bonding pattern and dynamics of granisetron during MD simulations. **a**. The inset shows the distributions of the dihedral angle between the two moieties of granisetron (drawn in red), either bound to the receptor or free in water. The position of the aromatic part is relatively conserved in the protein indicating that the conformations sampled result from the conformational dynamics of the azabicyclo ring in the binding site. **b**. The distance distribution depicts the H-Bonds between the ring -NH group and the backbone carbonyl of W156 monitored over the MD trajectory for the 5 subunits. **c**. Two snapshots illustrate the main conformations sampled by granisetron.

**Figure SI8.**
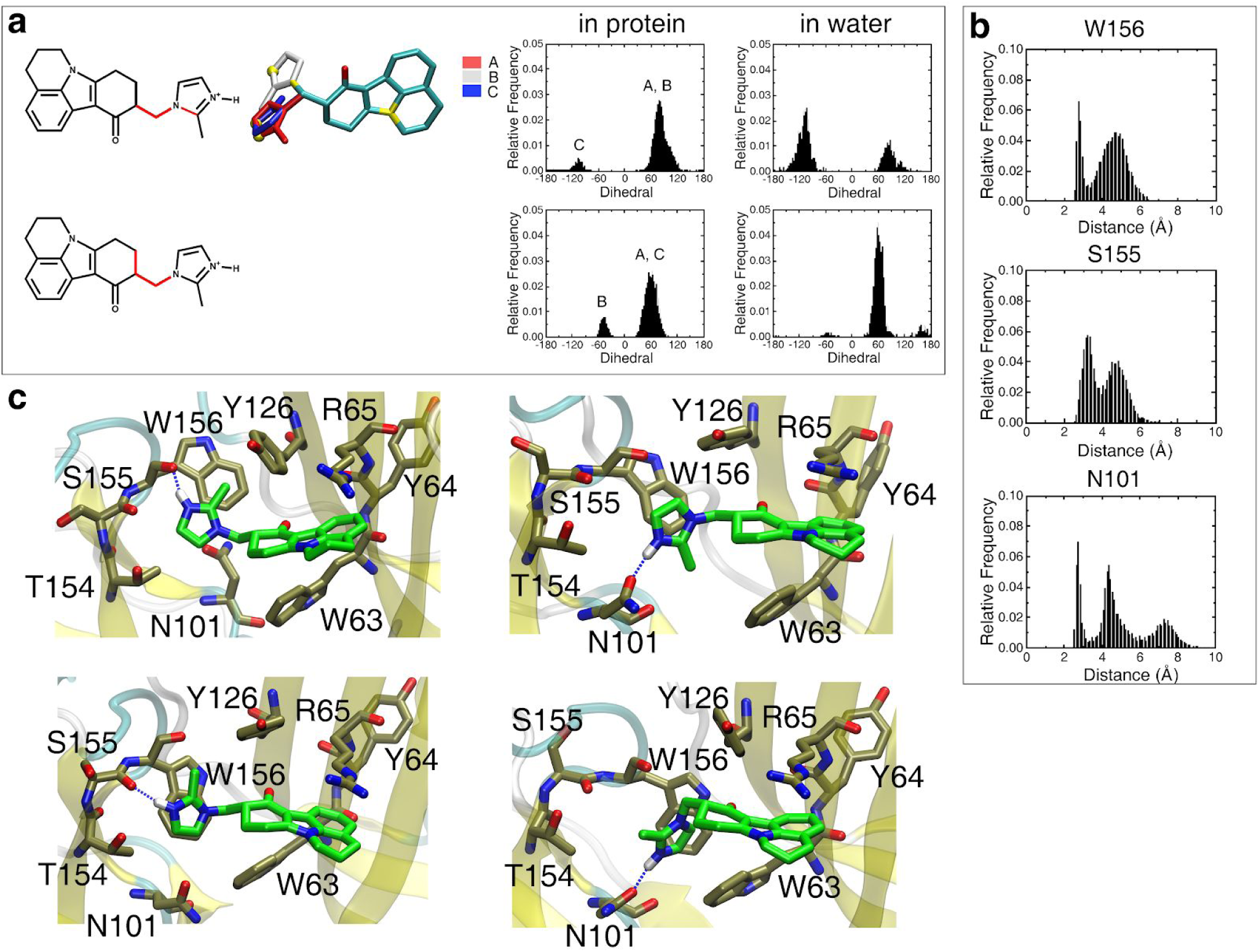
H-bonding pattern and dynamics of cilansetron during MD simulations. **a**. The inset shows the distributions of the two dihedral angles between the two moieties of cilansetron (drawn in red), either bound to the receptor or free in water. The position of the aromatic part is relatively conserved in the protein indicating that the conformations sampled result from the conformational dynamics of the azabicyclo ring in the binding site. **b**. The three distance distributions depict the H-Bonds between the ring -NH group and the backbone carbonyl of W156, the backbone carbonyl of S155 or the side chain carbonyl of N101 monitored over the MD trajectory for the 5 subunits. **c**. Four snapshots illustrate the main conformations sampled by cilansetron.

**Figure SI9.**
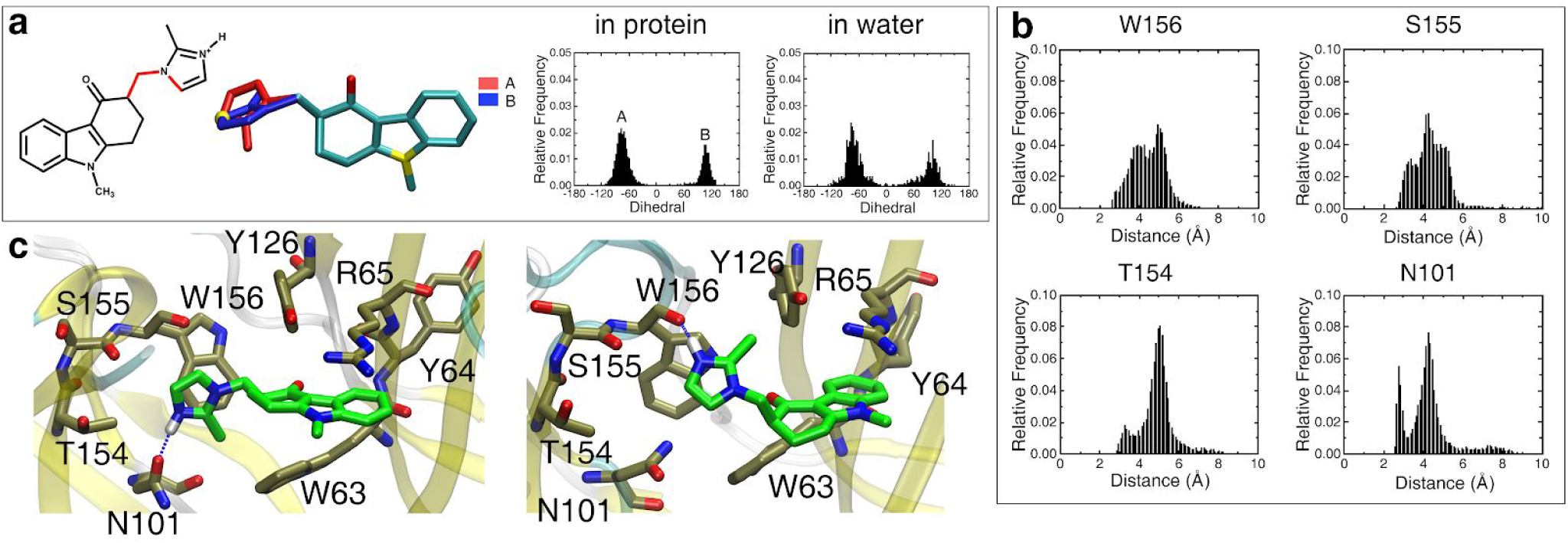
H-bonding pattern and dynamics of ondansetron during MD simulations. **a**. The inset shows the distributions of the dihedral angle between the two moieties of ondansetron (drawn in red), either bound to the receptor or free in water. The position of the aromatic part is relatively conserved in the protein indicating that the conformations sampled result from the conformational dynamics of the azabicyclo ring in the binding site. **b**. The four distance distribution depict the H-Bonds between the ring -NH group and the backbone carbonyl of W156, the backbone carbonyl of S155, the backbone carbonyl of T154 and the side chain carbonyl of N101 monitored over the MD trajectory for the 5 subunits. **c**. Four snapshots illustrate the main conformations sampled by ondansetron.

**Figure SI10.**
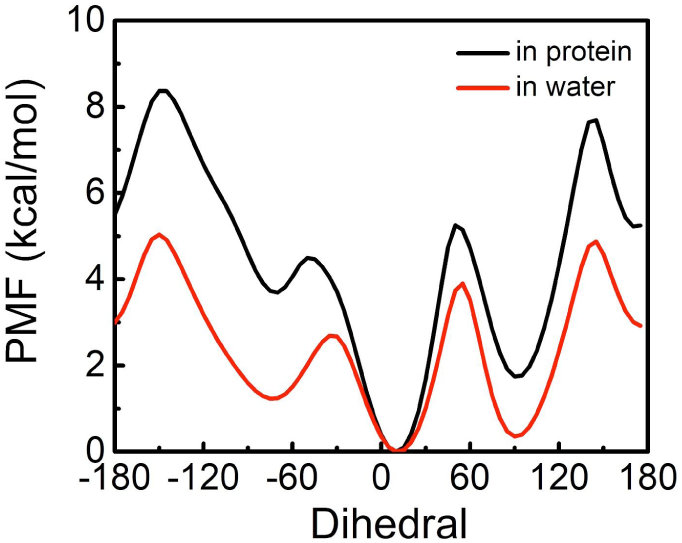
Potential of mean-force (PMF) associated with the palonosetron dihedral angle defined in Fig. 4. **a**. Free-energy calculations were performed using the e-ABF algorithm implemented in NAMD together with the colvar module. (black) PMF determined for the palonosetron in complex with 5-HT3 (200ns) (red) PMF determined for the palonosetron in water (400ns).

**Figure SI11.**
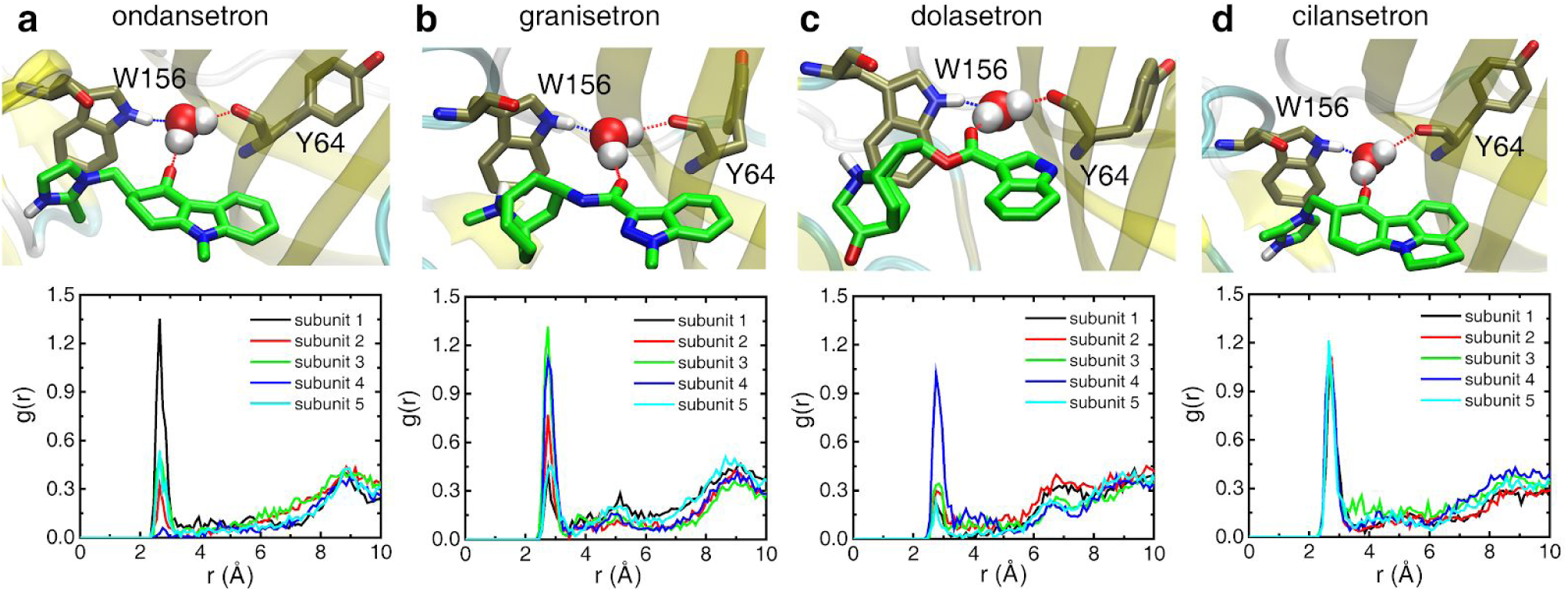
A trapped water molecule links (a) ondansetron, (b) granisetron, (c) dolasetron and (d) cilansetron H-bond acceptor and the receptor. **Upper panel**. Representative snapshot illustrating the water mediated H-Bond network linking the drug central carbonyl to protein residues W156 and Y64. **Lower panel**. Pair radial distribution functions between the oxygen of the drug central carbonyl and the water oxygen, characterizing the probability density to find the two species at a given distance. The peak located at ∼3 Å indicates that for every subunit, there is always one water molecule H-bonded to all drugs

**Figure SI12.**
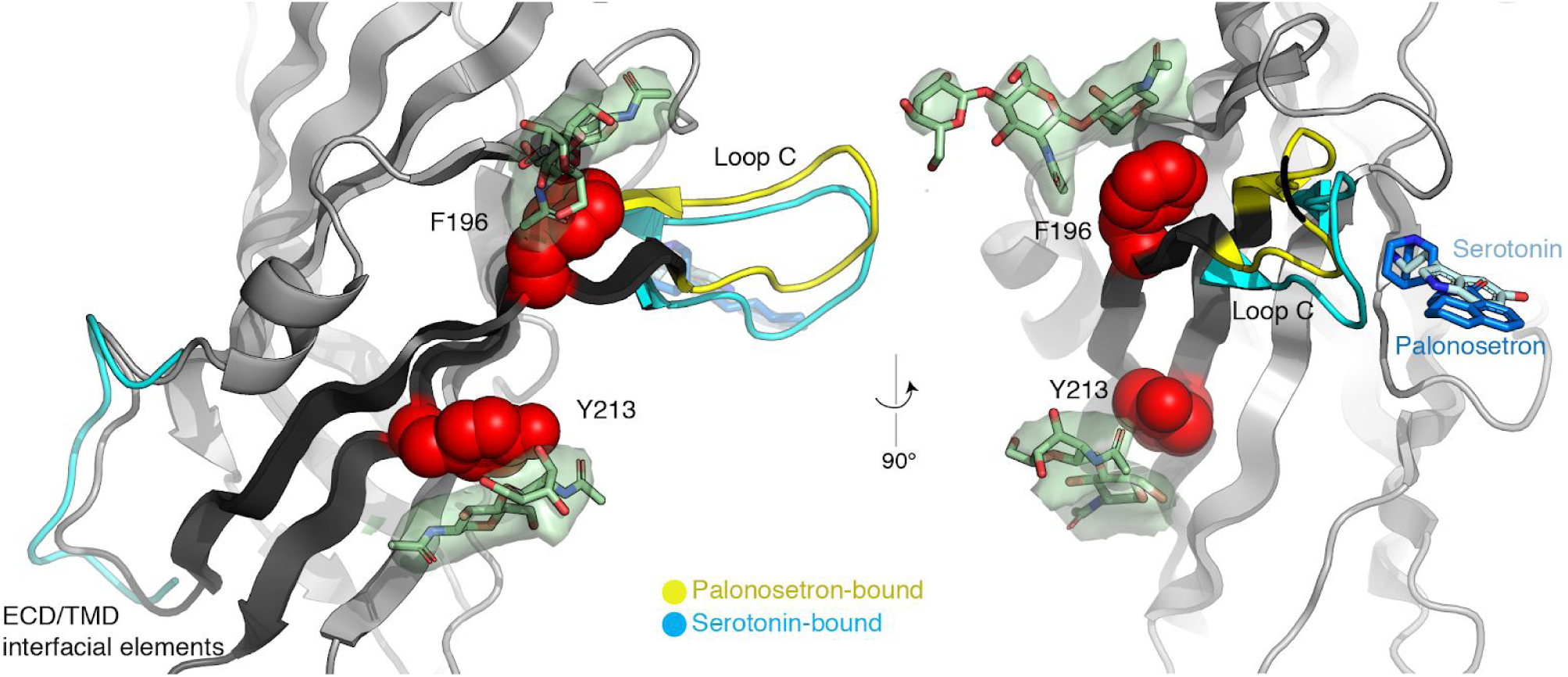
Carbohydrates clamp β9-β10. Two orthogonal cartoon representations of the principal subunit, with the loop C and the β8-β9 interfacial loop superimposed for the serotonin-bound active conformation (superimposition of the complementary subunit). N-linked carbohydrates are depicted in sticks and transparent surfaces for the Cryo-EM map, while residues F196 and Y213 of β9-β10 that interact with them are shown as red spheres. One may hypothesize that the clamp could help transmit loop C capping to the ECD/TMD interface.

**Figure SI13.**
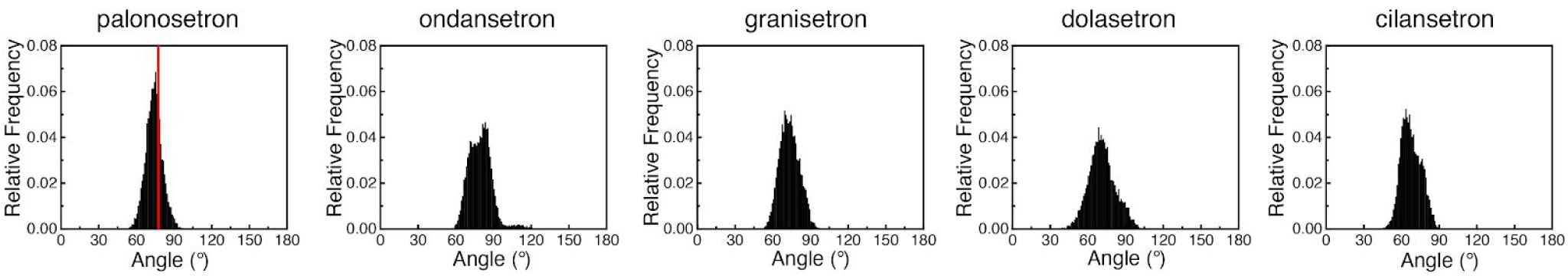
Distribution of the angle between the aromatic moiety and the pseudo-symmetry axis of the protein. The red line on the left panel corresponds to the value for the palonosetron cryo-EM structure.

**Table S2.** Cryo-EM data collection, refinement and validation statistics for the Palonosetron-bound 5-HT3 receptor, EMD-10673, PDB 6Y1Z.

| Data collection and processing |  |
| --- | --- |
| Magnification | 130'000 |
| Voltage (kV) | 300 |
| Electron exposure (e<sup>-</sup>/Å<sup>2</sup>) | 53 |
| Defocus range (μm) | -0.8 to -2.5 |
| Pixel size (Å) | 1.067 |
| Symmetry imposed | C5 |
| Initial particle images (no.) | 514'057 |
| Final particle images (no.) | 127'971 |
| Map resolution (Å)<br>FSC threshold | 2.8<br>0.143 |

| Refinement |  |
| --- | --- |
| Initial model used (PDB code) | 4PIR |
| Model resolution (Å)<br>FSC threshold | 3.13<br>0.5 |
| Model composition |  |
| Non-hydrogen atoms | 16'610 |
| Protein | 16'095 |
| Palonosetron | 110 |
| Carbohydrates | 405 |
| B factors (Å<sup>2</sup>) min/max/mean |  |
| Protein | 60.17/158.56/96.97 |
| Ligand | 76.35/133.10/104.23 |
| R.m.s. deviations |  |
| Bond lengths (Å) | 0.003 |
| Bond angles (°) | 0.537 |
| Validation |  |
| MolProbity score | 1.56 |
| Clashscore | 3.46 |
| Poor rotamers (%) | 2.21 |
| Ramachandran plot |  |
| Favored (%) | 97.11 |
| Allowed (%) | 2.89 |
| Disallowed (%) | 0 |

**Figure SI14.**
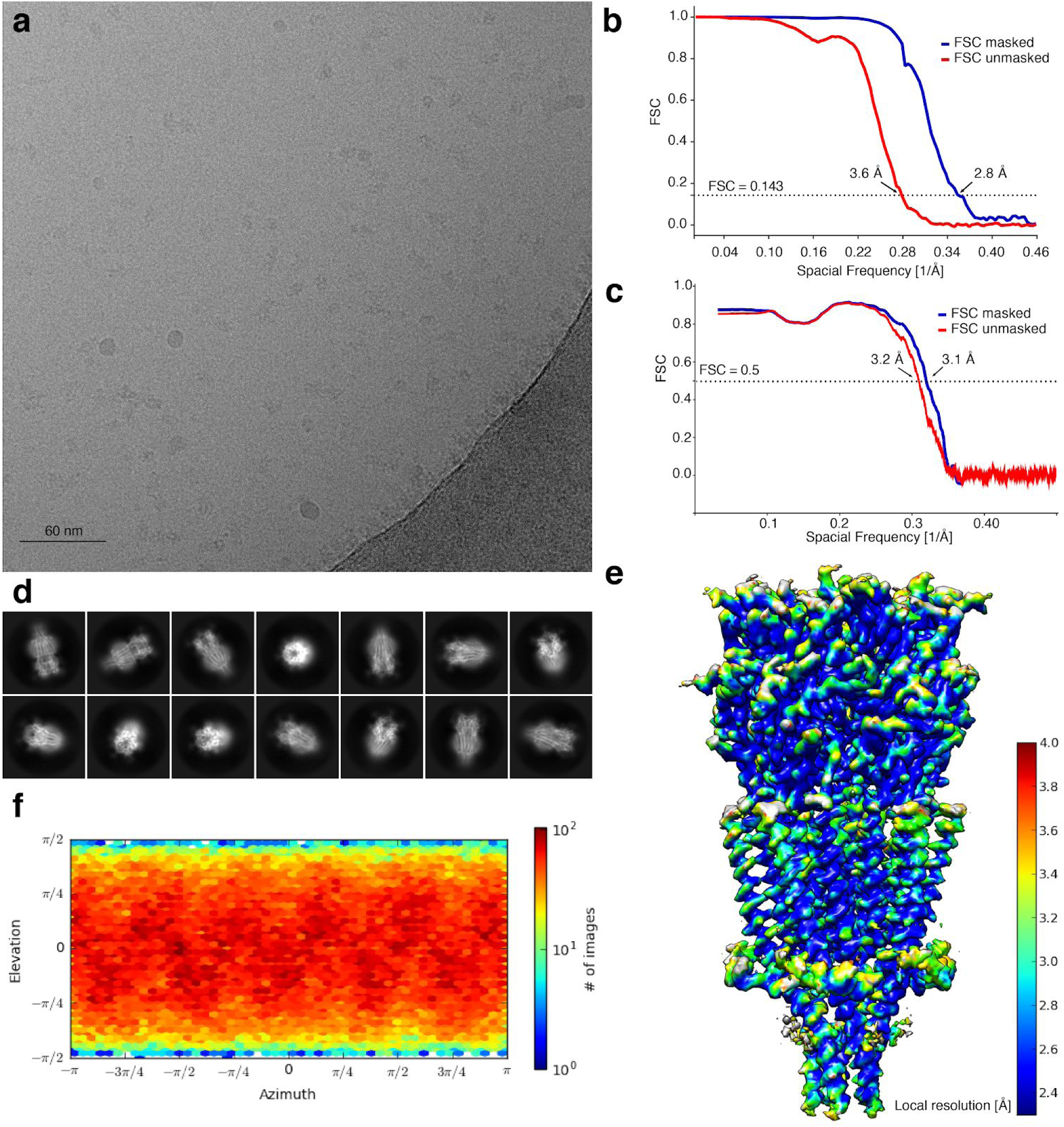
Image analysis workflow. **a**. A representative micrograph (out of 3037 micrographs) of the pre-incubated with palonosetron 5HT3A receptor in vitreous ice. **b**. Gold-standard Fourier shell correlation (FSC), calculated by comparing the two independently determined half maps in the final Non-Uniform refinement in cryoSPARC. The dotted line represents the 0.143 FSC threshold, which indicates a nominal resolution of 3.6 Å for the unmasked (red) and 2.8 Å for the masked (blue) reconstruction. **c**. Model-to-map Fourier shell correlation (FSC). The dotted line represents the 0.5 FSC threshold, which indicates a nominal resolution of 3.23 Å for the unmasked (red) and 3.13 Å for the masked (blue) reconstruction.**d**. The selected 2D class averages of the particles that were used for the final reconstruction. **e**. Side view of the unfiltered and unsharpened 3D density map colored according to the local resolution estimated in ResMap. **f**. Heat map of the angular distribution of particle projections for the final reconstruction calculated in cryoSPARC and colored-coded depending the approximate number of particles for each orientation.

**Figure SI15.**
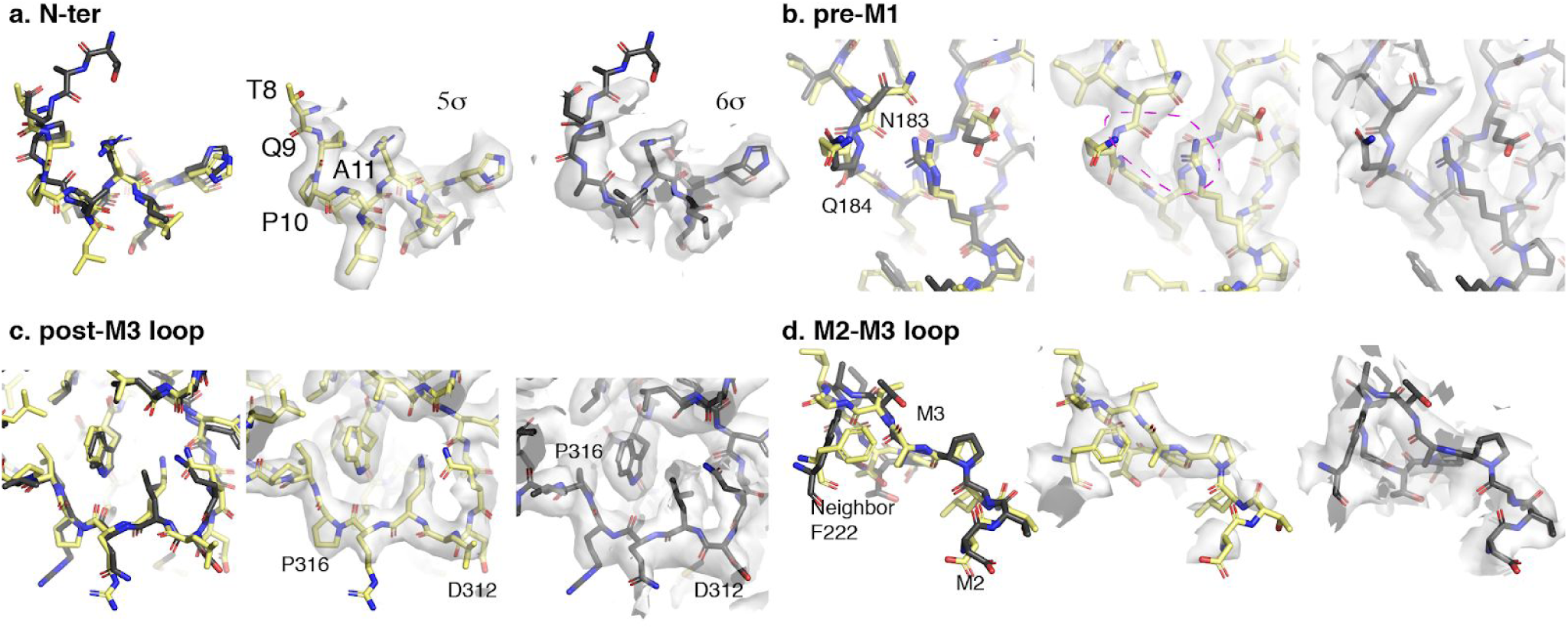
Comparison of the palonosetron- and granisetron-bound structures. In each panel, the palonosetron-bound cryo-EM structure reported here is superimposed to the granisetron-bound one obtained by Basak et al^16^; two views of structures in their density maps are provided. Globally the structures are highly similar, and only the few regions of divergence are shown here. These regions reflect zones of higher flexibility where the map can be interpreted in different models. **a**. N-terminal of the receptor. An offset before A11 exists. **b**. Pre-M1 region. Different orientations of the backbone of residues 183-184. Our conformation would favor the interaction of R219 with loop β8-β9. **c**. Post-M3 loop. This small loop was incorrectly modeled in former lower-resolution structures and we can now model D312 in a position where it will interact with the triplet of arginines that define single-channel conductance^79^. Differences here may influence putative mutagenesis work performed by others research groups. **d**. M2-M3 loop. This loop is flexible in almost any reconstruction we obtained so far. Despite the importance of this loop, it remains very hard to model.

## Footnotes

1 Throughout the report, mutants are simply noted *residue-number-residue* to facilitate the reading, without specifying if they are mutants of the human or of the mouse 5-HT3 receptor (as long as both receptors have the same amino-acid at a given position of course).

